# Fatty acid oxidation impairs macrophage effector functions that control *Mycobacterium tuberculosis*

**DOI:** 10.1101/799619

**Authors:** Pallavi Chandra, Li He, Matthew Zimmerman, Guozhe Yang, Stefan Köster, Mireille Ouimet, Han Wang, Kathyrn J. Moore, Véronique Dartois, Joel D. Schilling, Jennifer A. Philips

## Abstract

Macrophage activation involves metabolic reprogramming to support antimicrobial cellular functions. How these metabolic shifts influence the outcome of infection by intracellular pathogens remains incompletely understood. *M. tuberculosis* (Mtb) modulates host metabolic pathways and utilizes host nutrients, including cholesterol and fatty acids, to survive within macrophages. We found that intracellular growth of Mtb depends on host fatty acid catabolism: when host fatty acid β-oxidation (FAO) was blocked chemically with trimetazidine, a compound in clinical use, or genetically by deletion of the mitochondrial fatty acid transporter carnitine palmitoyltransferase 2 (CPT2), Mtb failed to grow in macrophages and its growth was attenuated in mice. Global metabolic profiling and mechanistic studies support a model in which inhibition of FAO generates mitochondrial reactive oxygen species, which enhance macrophage NADPH oxidase and xenophagy activity to better control Mtb infection. Thus, FAO inhibition promotes key antimicrobial functions of macrophages and overcomes immune evasion mechanisms of Mtb.

## INTRODUCTION

Macrophages are at the forefront of innate immune defense and are required for microbial killing, tissue homeostasis and repair, inflammation, and development. A wealth of scientific studies describes how these cells recognize pathogen-associated molecular patterns (PAMPs), which activate downstream signaling cascades to drive pro-inflammatory responses and resolve infection. Macrophage activation involves profound metabolic reprogramming to support specific phenotypic functions. Exposure of macrophages to stimuli such as IFN-γ and LPS induces an inflammatory phenotype characterized by enhanced glycolytic metabolism and impaired oxidative phosphorylation (OXPHOS), similar to the Warburg effect described for cancer cells. Glucose catabolism provides a rapid means to generate ATP, and boosts the pentose phosphate pathway (PPP) and tricarboxylic acid (TCA) cycle for the generation of important immuno-metabolites such as NADPH, itaconate, and prostaglandins (Galván-Peña and O’Neill, 2014; Kelly and O’Neill, 2015; Van den Bossche et al., 2017). On the other hand, cytokines such as IL-4 induce an anti-inflammatory phenotype in macrophages important for tissue homeostasis and anti-parasitic responses. These alternatively activated macrophages have increased FAO and OXPHOS (Galván-Peña and O’Neill, 2014). The majority of studies on macrophage metabolism focus on IFNγ/LPS-and IL-4-induced states, although accumulating evidence suggests that macrophage polarization states are multi-dimensional (Xue et al., 2014). The metabolic characterization of more diverse macrophage phenotypes, and their impact on antimicrobial capacity in the context of specific infections is largely unexplored.

Recent studies also highlight a link between *Mycobacterium tuberculosis* (Mtb) pathogenesis and host metabolism. Mtb is the causative agent of tuberculosis (TB), which kills more people yearly than any other infection. Macrophages are a main cellular niche of Mtb (Cohen et al., 2018; Wolf et al., 2007). Within the lung, alveolar (AM) and interstitial (IM) macrophages are the major populations of infected macrophages, and they have distinct metabolic profiles. AMs, which preferentially utilize FAO, represent a permissive niche for Mtb replication, whereas glycolytically active IMs restrict infection (Huang et al., 2018). Mtb alters macrophage metabolism along with shifting macrophage phenotype to a more pro-inflammatory state (Arts et al., 2016; Cumming et al., 2018; Gleeson et al., 2016; Ouimet et al., 2016; Shi et al., 2015). Mtb enhances the dependency of mitochondrial oxidative metabolism on fatty acids, in particular exogenous fatty acids, and induces the formation of lipid-droplet-filled or “foamy” macrophages (Cumming et al., 2018; Peyron et al., 2008a; Russell et al., 2009; Singh et al., 2012). Foamy macrophages are found within the inner layers of granulomas, a common histopathologic feature of TB, and the bacilli themselves can be found in close approximation to intracellular lipid droplets. It is thought that lipid bodies serve as a source of nutrients in the form of cholesterol esters and fatty acids for the bacilli, thus providing a hospitable niche for the bacterium (Brzostek et al., 2009; Daniel et al., 2011; Marrero et al., 2010; Munoz-Elias and McKinney, 2005; Pandey and Sassetti, 2008; Peyron et al., 2008b; Singh et al., 2012). Moreover, a growing body of literature suggests a link between lipid metabolism and cellular control of Mtb. A number of host-directed therapies (HDTs) under investigation for TB, such as statins and metformin, modulate host lipid metabolism (Parihar et al., 2014; Singhal et al., 2014). However, how these metabolic shifts influence the outcome of Mtb infection remains incompletely understood.

Cellular regulators of lipid metabolism such as microRNA-33 (miR-33) and the transcription factors PPAR-α and PPAR-γ play a role in the formation of Mtb-induced lipid droplets. Studies in which miR-33, PPAR-α, and PPAR-γ were modulated revealed a correlation between cellular lipids and intracellular survival of mycobacteria, such that increased intracellular lipids were associated with enhanced bacterial replication (Almeida et al., 2009; Kim et al., 2017; Ouimet et al., 2016). These studies led to the idea that impaired host fatty acid catabolism and the formation of foamy macrophages serves to enhance intracellular survival of Mtb, but direct evidence is lacking. Indeed, miR-33, PPAR-α, and PPAR-γ impact diverse aspects of host biology that influence the antimicrobial capacity of macrophages, including mitochondrial function and autophagy, and a causal link between the lipid changes they induce and the survival benefit to the bacilli has not been clearly established. Previously, we showed that induction of miR-33/33* in response to Mtb enhances intracellular survival of Mtb (Ouimet et al., 2016). We showed that this pro-bacterial function of miR-33/33* was related in part to its ability to block autophagy, and we speculated that its ability to block fatty acid catabolism and promote the formation of lipid droplets also enhanced bacterial replication. Here, we tested whether blocking fatty acid catabolism indeed promotes Mtb replication, as prevailing models predict. Instead, we found that inhibiting macrophage FAO chemically or genetically restricted intracellular growth of Mtb. FAO inhibition promoted a pathway of bacterial killing in which induction of mitochondrial ROS (mitoROS) lead to NADPH oxidase and autophagy-dependent control of Mtb.

## RESULTS

### Inhibiting FAO restricts growth of intracellular Mtb

miR-33 inhibits FAO by targeting genes such as carnitine palmitoyltransferase 1 (CPT-1) and the hydroxyacyl-CoA dehydrogenase trifunctional multienzyme complex subunit beta (HADHB). CPT-1 is required for the entry of long chain fatty acids into the mitochondrial matrix, while HADHB catalyzes the final step of *β*-oxidation. Small molecule inhibitors, etomoxir (ETM), oxfenicine (OXF), and trimetazidine (TMZ) also target these steps in FAO. ETM and OXF inhibit CPT-1, whereas TMZ blocks the 3-ketoacyl-CoA thiolase activity of HADHB (Figure 1A). Thus, to determine whether the inhibition of FAO conferred by miR-33 contributed to its ability to enhance the intracellular survival of Mtb, we tested the effect of chemical inhibition of FAO on intracellular bacterial replication. We infected murine bone marrow-derived macrophages (BMDMs) with the H37Rv strain of Mtb, and four hours later, we supplemented the media with ETM, OXF, or TMZ. After treatment, we estimated intracellular Mtb growth by plating for colony forming units (CFU) at 72 hours post infection (hpi). Unexpectedly, treatment with all three FAO inhibitors restricted the intracellular growth of Mtb compared to untreated controls. We observed a dose-dependent reduction in Mtb CFUs in macrophages treated with micromolar concentrations of ETM (Figure 1B), while TMZ and OXF were effective at nanomolar concentrations (Figures 1C and D). The antitubercular activity of TMZ was corroborated using a live-dead reporter strain of Mtb (Figure S1A). By comparison, metformin (MET), which has previously been shown to restrict intracellular Mtb, worked at millimolar concentrations (Singhal et al., 2014) (Figures 1B-D). Using calcein fluorescent dye, we found that macrophage viability was unaffected by FAO inhibitors ETM and TMZ (Figure S1B). In addition, the inhibitors did not have direct toxicity on Mtb in broth culture (Figure S1C and S1D), and they enhanced macrophage control against a distantly related mycobacterium, *Mycobacterium abscessus* (Figure S1E). Consistent with the ability to impair FAO, TMZ enhanced cellular lipid accumulation based upon BODIPY staining (Figure S1F). In addition, in keeping with impaired fatty acid oxidation, TMZ treated macrophages had reduced oxygen consumption (Figure S1G), similar to macrophages from mice that were genetically deficient in FAO due to deletion of carnitine palmitoytransferase 2 (*Cpt2^fl/fl^* LysM-Cre^+^*; Cpt2* cKO) (Gonzalez-Hurtado et al., 2017).

**Figure 1.**
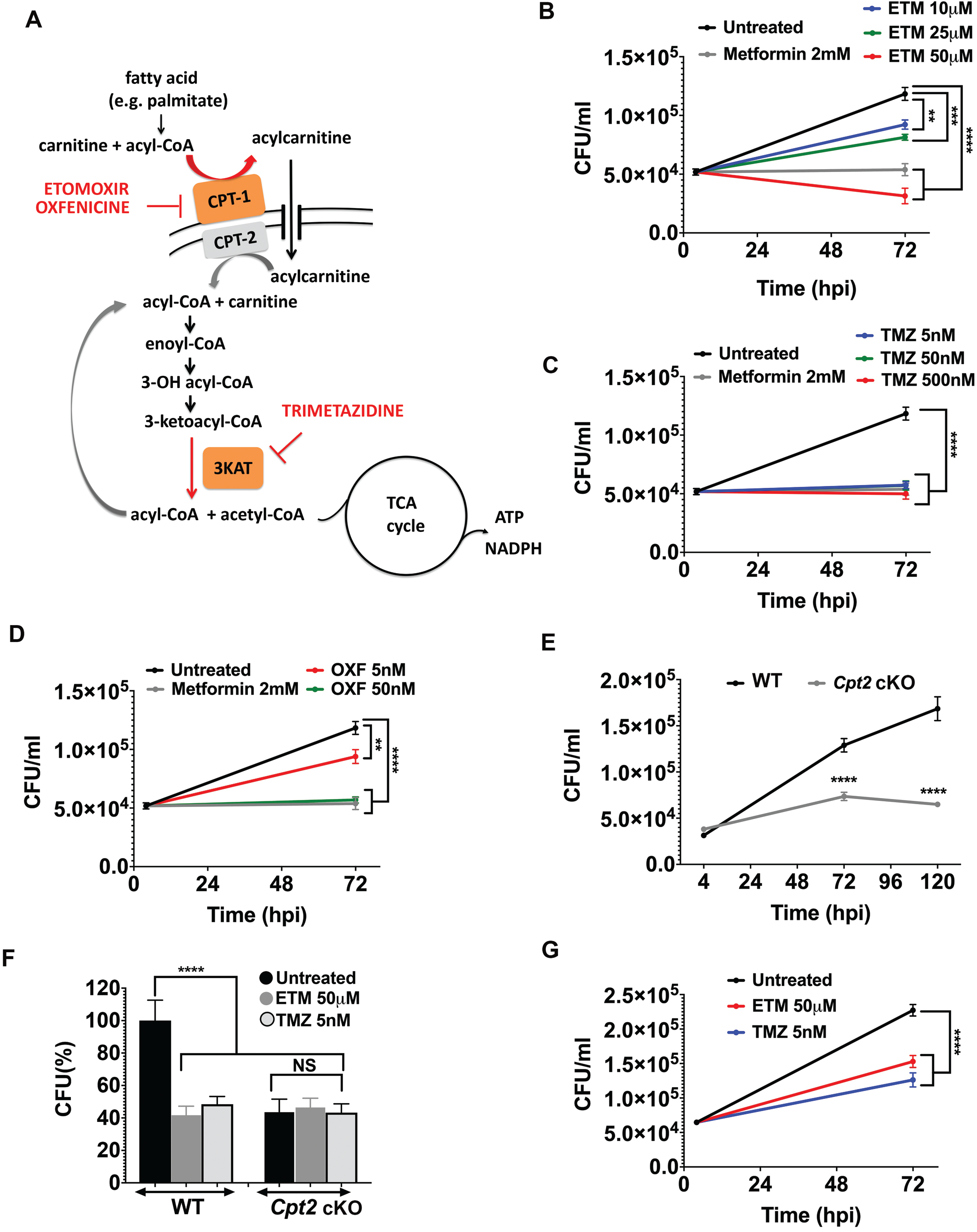
Inhibition of macrophage FAO restricts intracellular Mtb growth. **(A)** In the carnitine shuttle, long chain fatty acids that have been activated to acyl-CoA derivatives are converted to acylcarnitines by CPT1 at the outer mitochondrial membrane. Etomoxir and oxfenicine inhibit CPT1. Acylcarnitines are transported across the mitochondrial membrane by a dedicated translocase. In the mitochondria, CPT2 converts acylcarnitines back to acyl-CoA and carnitine. Acyl-CoA chains then undergo β-oxidation, successively generating acetyl-CoA that enters the TCA cycle. Trimetazidine inhibits 3-ketoacyl-CoA thiolase, which catalyzes the release of acetyl-CoA from the acyl-CoA chain. **(B-D)** Survival of Mtb (H37Rv) in BMDMs that were untreated or treated for 72 h with 2 mM metformin (MET) or indicated concentrations of ETM **(B),** TMZ **(C),** or OXF **(D). (E)** Survival of Mtb in BMDMs from *Cpt2* KO (*Cpt2^fl/fl^*LysM-Cre^+^) and littermate controls (*Cpt2^fl/f^* Cre^-^, WT) 4, 72 and 120 hpi. **(F)** Survival of Mtb in BMDMs from *Cpt2* cKO and WT controls treated with ETM, TMZ, or vehicle control for 72h. CFU are expressed as % of WT untreated control. **(G)** PMA-differentiated THP-1 cells were infected with Mtb and treated with ETM or TMZ for 72 hpi prior to enumerating CFU. (**B**-**G**) Data shows mean +/-s.e.m. from one representative experiment from at least 2 independent experiments.**P ≤ 0.009, ***P=0.0003, ****P<0.0001, NS-not significant, one-way ANOVA (**B**-**D**, **F**-**G**) or unpaired Students t-test with Welch’s correction (**E**). See also Figure S1.

To confirm that the anti-mycobacterial activity was indeed a result of FAO inhibition, we compared the intracellular growth of Mtb in *Cpt2* cKO macrophages to littermate controls. As shown in Figure 1E, Mtb was significantly impaired in *Cpt2* cKO macrophages as compared to control. Moreover, FAO inhibitors lacked activity in *Cpt2* cKO macrophages, confirming their target specificity (Figure 1F). To determine whether FAO inhibitors were also active against Mtb in human macrophages, we treated PMA-differentiated THP-1 cells. As we found in murine macrophages, FAO inhibitors impaired Mtb growth in THP-1 cells (Figure 1G). Taken together, our findings suggest that FAO inhibition enhances the ability of macrophages to control Mtb infection. Since ETM is documented to have off-target effects (Divakaruni et al., 2018) and TMZ was effective at nanomolar concentrations, we elected it for further study.

### Inhibiting FAO restricts growth of Mtb *in vivo*

Given our *in vitro* findings, we investigated whether inhibition of FAO impacts mycobacterial control in mice. We compared Mtb infection in mice with myeloid-specific knockout of *Cpt2* (*Cpt2^fl/fl^* LysM-Cre^+^*; Cpt2* cKO) and littermate controls. We exposed mice to Mtb by aerosol and estimated lung bacterial burden after 7 days. We observed that growth of Mtb was significantly reduced in *Cpt2* cKO mice compared to controls (Figure 2A). To determine whether chemical inhibition of FAO had antimicrobial activity in mice, we first performed pharmacokinetic (PK) studies to establish a TMZ dosing strategy (Figure S2A and S2B). The half-life of TMZ in people is approximately 6 hours, but in mice the half-life by oral or IV route was less than 1 hour, so we used Alzet osmotic pumps to deliver a continuous dose over 2 weeks (Figure S2C). First, we examined the ability of TMZ to reduce bacterial burden during acute infection. In the first study we infected mice with 1000-2000 Mtb CFUs and treated them with saline or TMZ for 2 weeks. We found that mice treated with 16.8 mg/kg/day TMZ experienced a 67% reduction in lung bacterial burden compared to animals in the control group (Figure 2B). A significant decrease in spleen CFU was also observed with 16.8 mg/kg/day TMZ treatment (Figure 2C). In a second study, we infected mice with 3000 CFUs, tested a higher TMZ dose range, and included PK analysis of infected animals. In this study, 16.8 mg/kg/day showed a trend towards reduced bacterial growth, which was not statistically significant, while the animals receiving 50 mg/kg/day experienced a 29% reduction in bacterial burden compared to saline controls (Figure 2D **and** 2E). In this group, TMZ treatment did not reduce Mtb CFU in spleen. During acute infection, the majority of Mtb bacilli are found within alveolar macrophages, suggesting that TMZ was active in this cell population. As infection progresses, additional phagocytic cells, including recruited monocytes, neutrophils, and dendritic cells become increasingly infected (Cohen et al., 2018; Wolf et al., 2007). To test whether TMZ was active in the setting of chronic infection, we began TMZ treatment 5 weeks post-infection and analyzed bacterial burden after 2 weeks of drug treatment. PK analysis was performed on infected mice 5 days after start of therapy and at end of treatment. In the setting of chronic infection, we found that TMZ treatment reduced bacterial burdens in the lungs by 35%, and in the spleen by 23% and 43%, in the 16.8 mg/kg and 50 mg/kg groups (Figure 2F **and** 2G). TMZ serum concentrations of 130 ng/ml and 370 ng/ml were achieved at the end of therapy for the 16.8 and 50 mg/kg/day groups, respectively (Figure 2H). TMZ is approved by the European Medicines Agency to treat angina, and the C_max_ obtained after a single 40 mg oral dose in humans is 127 ng/ml. Because there is higher serum protein binding in mice than humans (Figure S2D), the level achieving a similar free drug concentration in mice would be approximately 164 ng/ml. Thus, TMZ showed antimycobacterial activity at drug concentrations that are used clinically. In addition, consistent with the extremely wide therapeutic index of TMZ in preclinical toxicology studies (Harpey et al., 1988), no treatment related morbidity or mortality was observed in any of the studies. Overall, we conclude that inhibiting host fatty acid metabolism restricts mycobacterial growth *in vivo*.

**Figure 2.**
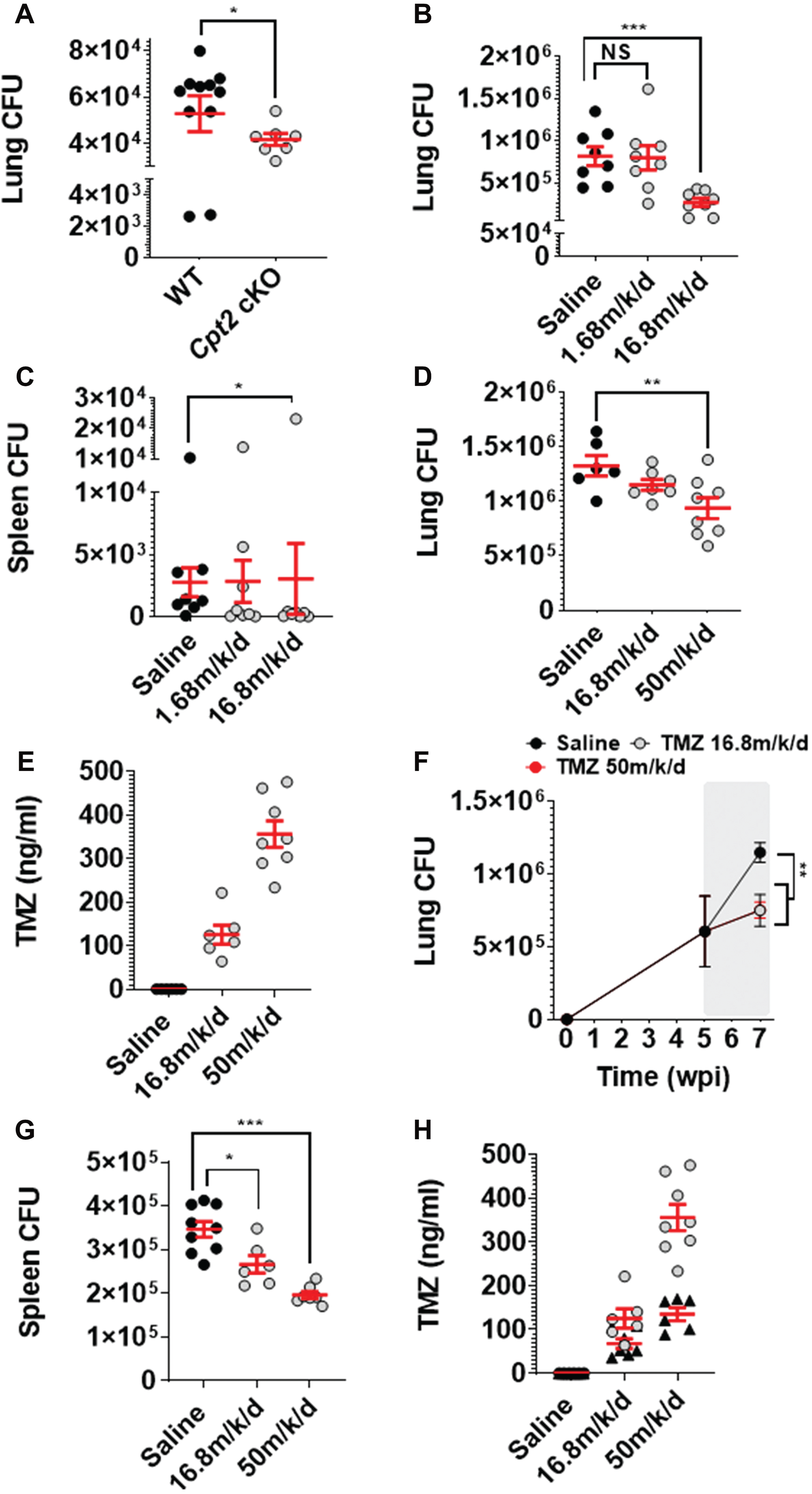
FAO inhibition restricts growth of Mtb *in vivo*. **(A)** *Cpt2* cKO mice and littermate controls were infected by aerosol with 100-200 Mtb CFU per animal, and CFU were enumerated from lungs 7 days post-infection. *P=0.04, **(B-D)** Efficacy of TMZ treatment was tested in mice that were acutely infected with Mtb. Plots show total lung and spleen CFU for the first study (B, C) and total lung CFU for the second study (D) after 2 weeks of TMZ treatment. *P=0.03, **P=0.02, ***P=0.0002. **(E)** Serum levels of TMZ were measured at the end of therapy in mice shown in (D). **(F,G)** In the chronic study, mice that were infected by aerosol with Mtb for five weeks were treated with TMZ or saline control for 2 weeks (shown in gray region) and total lung and spleen CFU were enumerated. *P=0.01, **P≤0.005, ***P=0.0002. **(H)** Serum levels of TMZ were measured in mice shown in (F) after 5 days of starting treatment (▴) and at the end of therapy (○). Data shows mean +/-s.e.m., P-values calculated using Mann-Whitney rank sum test. See also Figure S2.

### FAO inhibition enhances metabolic changes in macrophages in response to Mtb infection

We turned to *in vitro* studies to assess how host fatty acid metabolism was influencing control of Mtb. Given the increasing appreciation of a link between metabolism and immunity, we globally profiled metabolites to assess the effect of Mtb infection alone or when macrophage FAO was inhibited. Uninfected macrophages or those that had been infected for 4h were treated for 3h or 24h with TMZ or solvent control before profiling (Figure 3A). More than 500 metabolites were detected, and in agreement with previous reports, Mtb infection caused dramatic changes in macrophage metabolites (Figure 3: Figure S3; Table S1); we observed evidence of increased glycolytic and pentose phosphate pathway (PPP) flux (Figure 3B), itaconate production (Figure 3C), ROS and reactive nitrogen intermediates (RNI) production (Figure 3DE, 3E **and** Figure S3A), phospholipid and sphingomyelin changes (Figure S3B), and tryptophan metabolism (Figure S3C). As discussed below, on its own TMZ did not significantly alter metabolic profiles in uninfected macrophages, but in some cases it enhanced infection-induced changes.

**Figure 3.**
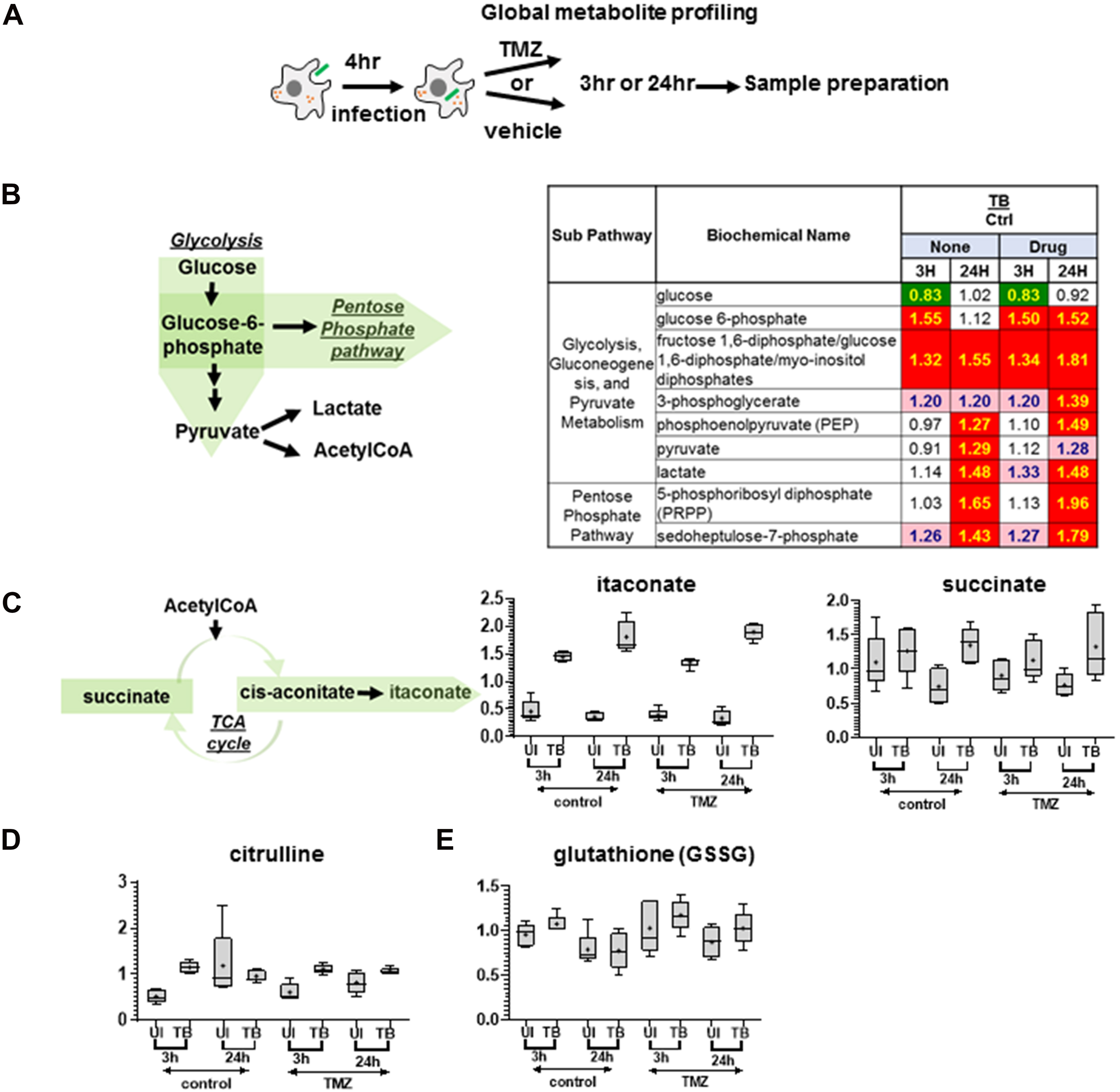
FAO inhibition enhances metabolic changes in response to Mtb infection. **(A)** Global metabolite profiling was performed on uninfected or Mtb infected BMDMs to assess the effect of infection alone or after treatment with 50nM TMZ for 3 or 24 h. **(B)** Macrophage glycolysis was augmented in response to Mtb infection, and a sharp accumulation in glycolytic intermediates was observed. Glucose-6-phosphate was shunted to the pentose phosphate pathway for the production of PRPP and sedoheptulose-6-phosphate. The pathway heat map shows metabolite ratios in Mtb infected and uninfected (TB/Ctrl) macrophages, that were untreated (None) or TMZ treated (Drug) for 3 or 24 h. Here, significant difference (p≤0.05, three-way ANOVA) between groups are colored in green for metabolite ratio of <1 and red for ratio of >1. Light green and red colors show groups that narrowly missed statistical cutoff for significance 0.05<p<0.10. **(C)** Mtb infection induced levels of intermediates itaconate and succinate. This reflected breakpoints in the TCA cycle that are characteristic of inflammatory macrophage phenotype. Innate immune response of macrophage against the pathogen manifested as elevations in **(D)** citrulline, an iNOS by product, and **(E)** oxidized glutathione, a host redox metabolite. (C-E) Graphs show ScaledImpData of different biochemicals in uninfected (UI) and infected (TB) samples at the indicated time points. The line and dot show data median and mean, respectively. See also Figure S3.

We observed increases in several glycolytic intermediates (Figure 3B), which suggest augmented glucose catabolism in Mtb-infected macrophages. Early after infection, glucose levels declined in Mtb versus control groups, while glucose-6-phosphate and hexose diphosphates (detected as isobar) accumulated. After 24h of Mtb infection, the glucose levels were comparable to uninfected controls, but we observed accumulation in glycolytic intermediates such as phosphoenolpyruvate. The magnitude of responses induced by infection appeared to be greater in TMZ-treated groups. For instance, 24 hpi glucose-6-phosphate was increased 1.52-fold increase by Mtb in drug-exposed cells and 1.12-fold in the absence of TMZ. Glucose can be shunted to the pentose phosphate pathway (PPP) resulting in formation of NADPH reducing power and pentose intermediates for nucleotide biosynthesis. We found that metabolites specific to the PPP, 5-phosphoribosyl diphosphate (PRPP) and sedoheptulose-7-phosphate, increased over time in Mtb-infected macrophages, and again, the level of these metabolites was slightly greater upon TMZ treatment (Figure 3B, Table S1).

Classically activated macrophages utilize TCA cycle intermediates for anabolic process and immune pathway signaling. They have a characteristic TCA cycle breakpoint in the conversion of isocitrate to α-ketoglutarate. This allows production of itaconate, an antimicrobial metabolite, from cis-aconitate (Figure 3C) (Lampropoulou et al., 2016; Michelucci et al., 2013; Nair et al., 2018). A second TCA cycle breakpoint occurs after succinate. As expected, we observed markedly increased itaconate in infected macrophages, as well as elevated succinate (Figure 3C). There was no substantial difference in the infection-induced elevation in itaconate or succinate in response to TMZ treatment, suggesting these metabolites did not account for the enhanced antimicrobial effect of TMZ.

The global metabolomics data also showed evidence of increased oxidative stress in Mtb infected in macrophages relative to uninfected controls, which appeared more pronounced in TMZ-treated samples compared to untreated controls. We observed elevations in citrulline levels early after infection, reflecting the activity of iNOS, which converts arginine to citrulline and NO (Figure 3D). This was accompanied by significant increases in dihydrobiopterin and biopterin (Figure S3A). These compounds represent oxidized forms of tetrahydrobiopterin, a cofactor in NO synthesis. Early after infection, the antioxidants glutathione and opthalmate, a compositional derivative of glutathione, were increased by infection, suggesting enhanced glutathione synthetase activity (Figure S3A). This was accompanied by increases in gamma-glutamyl amino acids, which reflect the transfer of gamma-glutamyl moiety of glutathione to acceptor amino acids. (e.g. gamma-glutamylglutamine) (Figure S3A). After 24 hpi, we observed increases in betaine, dimethylglycine, S-adenosylmethionine (SAM), S-adenosylhomocysteine (SAH) and cystathione (Figure S3A). These shifts could indicate increased methionine to cysteine conversion to support glutathione synthesis. Infected cells appeared to have a more oxidized intracellular milieu after TMZ treatment, as we observed more oxidized glutathione (Figure 3E). Additionally, at the later time point, there were greater infection-induced shifts in SAM, SAH and cystathione levels in infected TMZ-treated cells compared to untreated controls. Combined, the alterations in the glycolytic pathway, itaconate production, and redox homeostasis suggest that macrophages adopt a more M1-like metabolic phenotype in response to Mtb infection, and TMZ-induced perturbations may enhance or prolong these changes, but do not dramatically alter the response.

### FAO inhibition triggers mitochondrial ROS to promote mycobacterial control

The metabolomics data did not point to a particular metabolite or metabolic pathway to explain the antimicrobial activity of TMZ. Additionally, we measured the production of TNF-*α*, IL-6, CXCL2, IFN-*β*, CCL2, and CXCL10 in response to infection and TMZ treatment, and we did not observe a cytokine-driven antimicrobial response upon TMZ treatment (Figure S4). Since TMZ acts in the mitochondria, a major site of ROS production, and there was evidence of an increase in oxidative stress in TMZ-treated macrophages, we examined whether FAO inhibition promotes ROS production. We treated uninfected immortalized BMDMs (iBMDMs) with TMZ and observed an increase in ROS as early as 3 hours after treatment (Figure 4A). Addition of mitoTEMPO, a mitochondrial ROS (mitoROS) scavenger, abolished this ROS burst, whereas DPI, which inhibits NADPH oxidase had no effect, suggesting that the ROS was induced in the mitochondria (Figure 4A). Toll-like receptor activation and infection with Gram negative bacteria have previously been shown to result in mitoROS production (Mills et al., 2016; West et al., 2011), and Mtb infection induced a small amount of mitoROS (Figure 4B). However, significantly more ROS was generated in Mtb-infected BMDMs after TMZ treatment (Figure 4C). ROS levels in Mtb-infected *Cpt2* cKO BMDMs were also significantly higher than in control macrophages (Figure 4D). The TMZ-induced ROS still occurred in macrophages that lacked the NADPH oxidase (*Nox2* KO), consistent with a mitochondrial source (Figure 4C). We confirmed that TMZ induced mitoROS by using MitoSox fluorescent dye (Figure 4E). Combined these results demonstrate that TMZ induces ROS from a mitochondrial source after 3 hours of treatment in both infected and uninfected macrophages.

**Figure 4.**
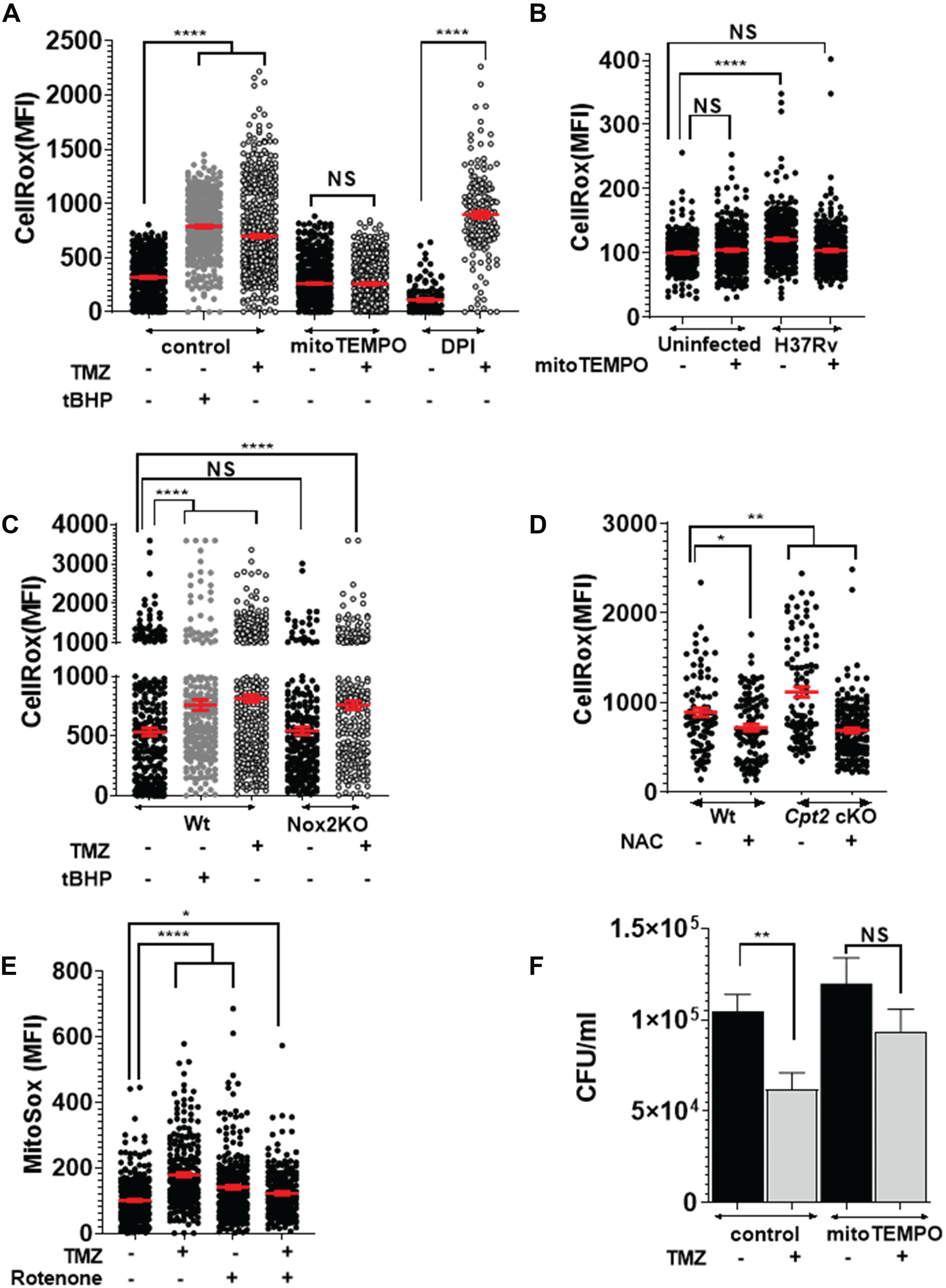
FAO inhibition induces mitochondrial ROS burst required for Mtb killing. CellRox mean fluorescence intensity (MFI) of **(A)** uninfected iBMDMs that were untreated or treated with 500nM TMZ for 3h, alone or in presence of mitoTEMPO (10µM) or DPI (10mM), **(B)** uninfected and infected BMDMs with or without mitoTEMPO (10µM) 4hpi, (C) Mtb-infected Wt and *Nox2* KO BMDMs that were untreated or treated with 5nM TMZ for 3h, and (D) Mtb-infected *Cpt2* cKO and control BMDMs 24 hpi that were untreated or treated with N-acetyl cysteine (NAC, 10mM). tert-Butyl hydroperoxide (tBHP, 0.25mM, 30 mins) was used as a positive control. *P=0.02, **P=0.001, ****P<0.0001, NS-not significant, calculated using unpaired Student’s t-test with Welch’s correction (A) and ordinary one-way ANOVA (B-D). (**E**) MitoSox MFI of uninfected BMDMs that were untreated or treated with 500nM TMZ for 3h, alone or in presence of rotenone (10µM, 30mins). *P=0.02, ****P≤0.0001, ordinary one-way ANOVA. **(F)** Mtb CFU in BMDMs that were untreated or treated with 50nM TMZ alone or in combination with mitoTEMPO for 120 hr. Data from one of 2 independent experiments shows mean +/- s.e.m.**P=0.005 calculated using unpaired Student’s t-test with Welch’s correction. **(A-E)** Data shown is mean +/- s.e.m. of MFI derived from individual cells in the sample. At least 100 cells were analyzed from duplicate samples. For (B, E), values are expressed as %uninfected. See also Figure S4.

We hypothesized that FAO inhibition resulted in mitoROS generation because of perturbed electron flow within the electron transport chain (ETC). ROS can be generated in multiple sites along the ETC during forward electron transport, as well as when electrons flow in reverse through complex I (NADH:coenzymeQ reductase) (Scialò et al., 2017). To determine the site within the ETC where TMZ induced ROS, we measured mitoROS production in macrophages treated with TMZ in combination with rotenone, an inhibitor of complex I. When electrons are flowing in the forward direction, rotenone prevents electron transport to CoQ, resulting in ROS generation. In contrast during reverse electron transport (RET), rotenone reduces ROS by preventing CoQ from transferring electrons back to complex I, where the RET-ROS is generated. We treated uninfected BMDMs with TMZ for 3h, and thirty minutes prior to measuring mitoROS we added rotenone. As expected, rotenone and TMZ on their own increased mitoROS production. In contrast, rotenone decreased the amount of ROS generated in response to TMZ treatment (Figure 4E) These findings are consistent with the idea that under conditions of TMZ treatment there is enhanced ROS generated due to RET at complex I.

Since recent studies have shown that mitoROS contributes to microbial control in macrophages (Abuaita et al., 2018; West et al., 2011), we asked whether TMZ-induced mitoROS was important for infection control or simply a by-product of metabolic perturbations. To address this, we estimated Mtb burden in macrophages treated with TMZ alone or in combination with mitoTEMPO. As shown in Figure 4F, mitoTEMPO on its own had little effect on the infection, but it partially reversed the antimicrobial effect of TMZ treatment. Overall, we conclude that FAO inhibition promotes mitoROS production from the ETC, which enhances macrophage control of Mtb infection.

### FAO inhibition-induced mitochondrial ROS drives NADPH oxidase recruitment to phagosomes

Previous studies suggest a link between mitoROS and NADPH oxidase that contributes to macrophage defense (Garaude et al., 2016). Indeed, while 3 h of TMZ treatment significantly increased mitoROS in both wt and *Nox2* KO BMDMs (Figure 4C), we observed that the anti-Mtb activity of FAO inhibitors required NADPH oxidase (Figure 5A). This suggested that the antimycobacterial activity in FAO-inhibited macrophages actually depended upon both a mitochondrial source and NADPH oxidase. Normally, pathogen associated molecular patterns (PAMPs) promote the recruitment of the NADPH oxidase to microbial phagosomes immediately after phagocytosis (Nunes et al., 2013). However, Mtb impairs the recruitment of the NADPH oxidase to the mycobacterial phagosome (Köster et al., 2017; Sun et al., 2013). Remarkably, at 24 hpi we observed that TMZ increased the recruitment of NADPH oxidase subunits gp91^phox^/NOX2 and p40^phox^ to the mycobacterial phagosomes (Figure 5B, C and D). Moreover, we found that NADPH oxidase recruitment was dependent on TMZ-induced mitoROS, as it was reversed with the addition of mitoTEMPO (Figure 5E). Thus, FAO inhibition appears to enhance two antimicrobial responses that are suboptimal during Mtb infection, mitoROS production and phagosomal recruitment of the NADPH oxidase. Mitochondria alone contributed to ROS within 3h of FAO inhibition, while the role of NADPH oxidase was appreciable at later time points, and macrophage control of Mtb depended upon both mitoROS and the NADPH oxidase.

**Figure 5.**
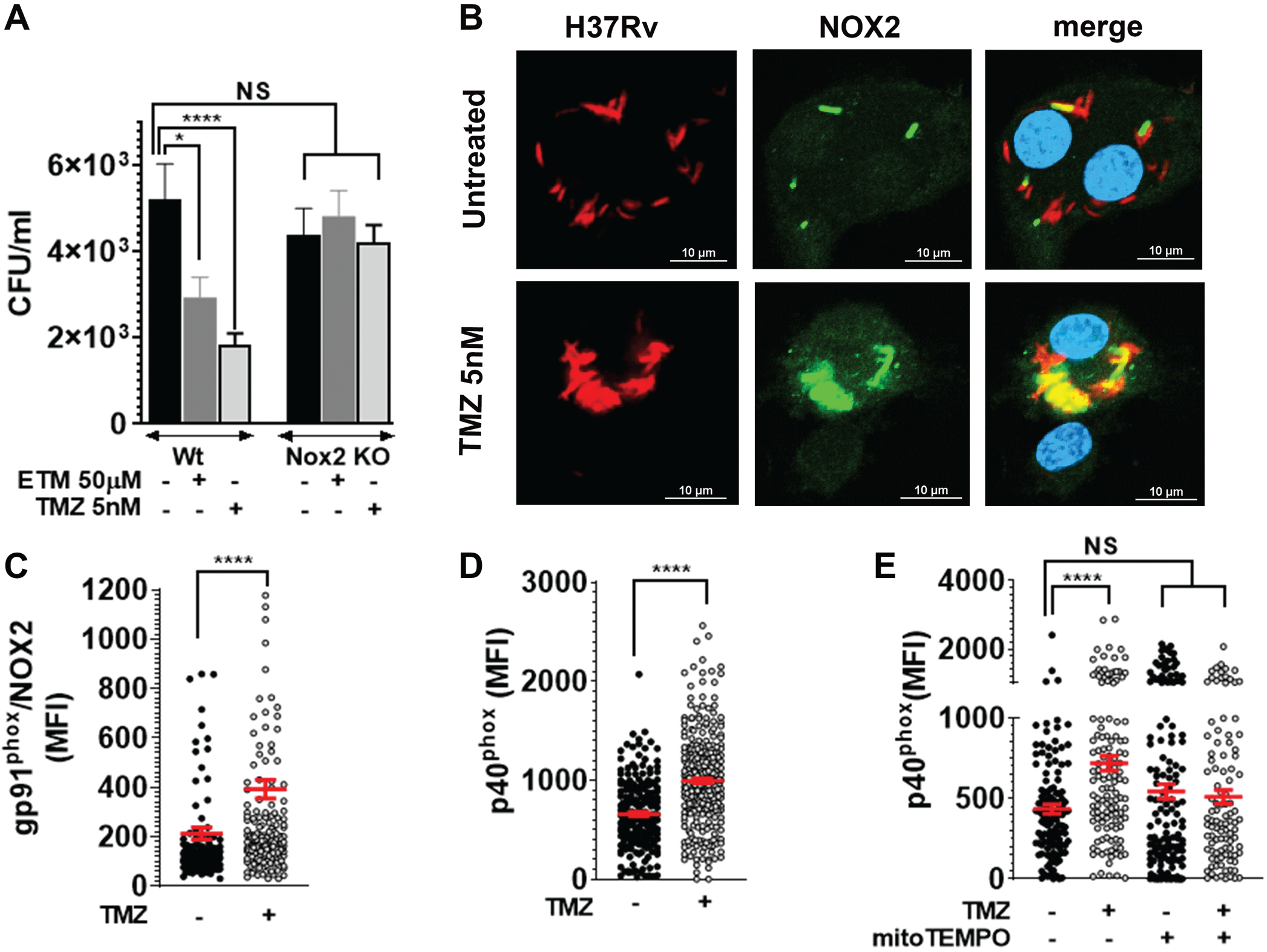
FAO inhibition promotes NADPH oxidase recruitment on phagosomes. **(A)** Survival of Mtb in BMDMs from *Nox2* KO and control mice that were untreated or treated with indicated concentrations of ETM and TMZ, 120 hpi.*P=0.04, ****P=0.0001, ordinary one-way ANOVA. **(B)** Immunofluorescence (IF) microscopy of gp91^phox^/NOX2 (green) and dsRed-expressing H37Rv (red) in BMDMs that were infected and treated with 5nM TMZ or untreated for 24h. The co-localized region is shown in yellow, scale bar = 10µm. MFI of the NADPH oxidase subunits **(C)** gp91^phox^/NOX2 and **(D)** p40^phox^ co-localized with H37Rv in BMDMs that were untreated or treated with 5nM TMZ (24hpi). **(E)** MFI of p40^phox^ co-localized with Mtb in BMDMs treated with 5nM TMZ alone or in presence of mitoTEMPO (48 hpi). (C-E) Automated image analysis was used to quantify gp91^phox^/NOX2 or p40^phox^ MFI co-localized with over 100 bacilli in at least 2 independent experiments. Data shows mean +/- s.e.m. of MFI from around individual bacilli in the sample. *P=0.02, ****P≤0.0001 calculated using unpaired Student’s t-test with Welch’s correction (C-D) or ordinary one-way ANOVA (E).

### FAO inhibition promotes xenophagy to restrict intracellular Mtb growth

TMZ treatment in Mtb-infected macrophages resulted in increases in phospholipids and sphingomyelin, which might alter phagosomal trafficking (Figure S3B). In addition, ROS generated by the NADPH oxidase promote a trafficking pathway called LC3-associated phagocytosis (LAP). Both LAP and autophagy of microbes (xenophagy) are characterized by the association of lipidated LC3 (LC3-II) with microbe-containing vacuoles (Martinez et al., 2015). LAP and xenophagy depend upon certain common ATG (autophagy related) proteins, and they also have unique requirements. To determine whether either LC3-trafficking pathway was involved in the antimicrobial activity of TMZ, we compared the efficacy of TMZ between control and *Atg5* cKO macrophages, which are deficient in both LAP and xenophagy. Indeed, the antimycobacterial control established by FAO inhibition was reversed in *Atg5*-deficient macrophages (Figure 6A). Although the baseline ROS staining was lower in *Atg5* cKO macrophages compared to control, ROS was still induced by TMZ in *Atg5* cKOs, suggesting that ATG5 acts downstream of mitoROS production (Figure 6B). To distinguish xenophagy from LAP, we examined *Atg14l* cKO and *Parkin2* KO macrophages, which are specifically involved in xenophagy. As we had seen in *Atg5* cKO macrophages, the antimycobacterial control established by FAO inhibition was reversed in *Atg14l* cKO and *Parkin2* KO as well (Figure 6C **and** 6D). In addition, in wt macrophages, TMZ treatment resulted in enhanced co-localization between Mtb and the autophagy adaptor, p62 (Figure 6E). The enhanced colocalization between Mtb and p62 occurred after 24 hours treatment and was dependent on mitoROS (Figure 6F **and** 6G). Combined, these data demonstrate that inhibition of FAO promotes xenophagy to control intracellular Mtb growth. Indeed, autophagosomal targeting of Mtb was also inherently higher in *Cpt2* cKO macrophages as compared to wt controls (Figure 6H **and** 6I). Moreover, TMZ treatment facilitated xenophagy of a mutant Mtb strain lacking *esxA* (Δ*esxA*), which is unable to perforate phagosomes and usually does not activate xenophagy (Figure 6J).

**Figure 6.**
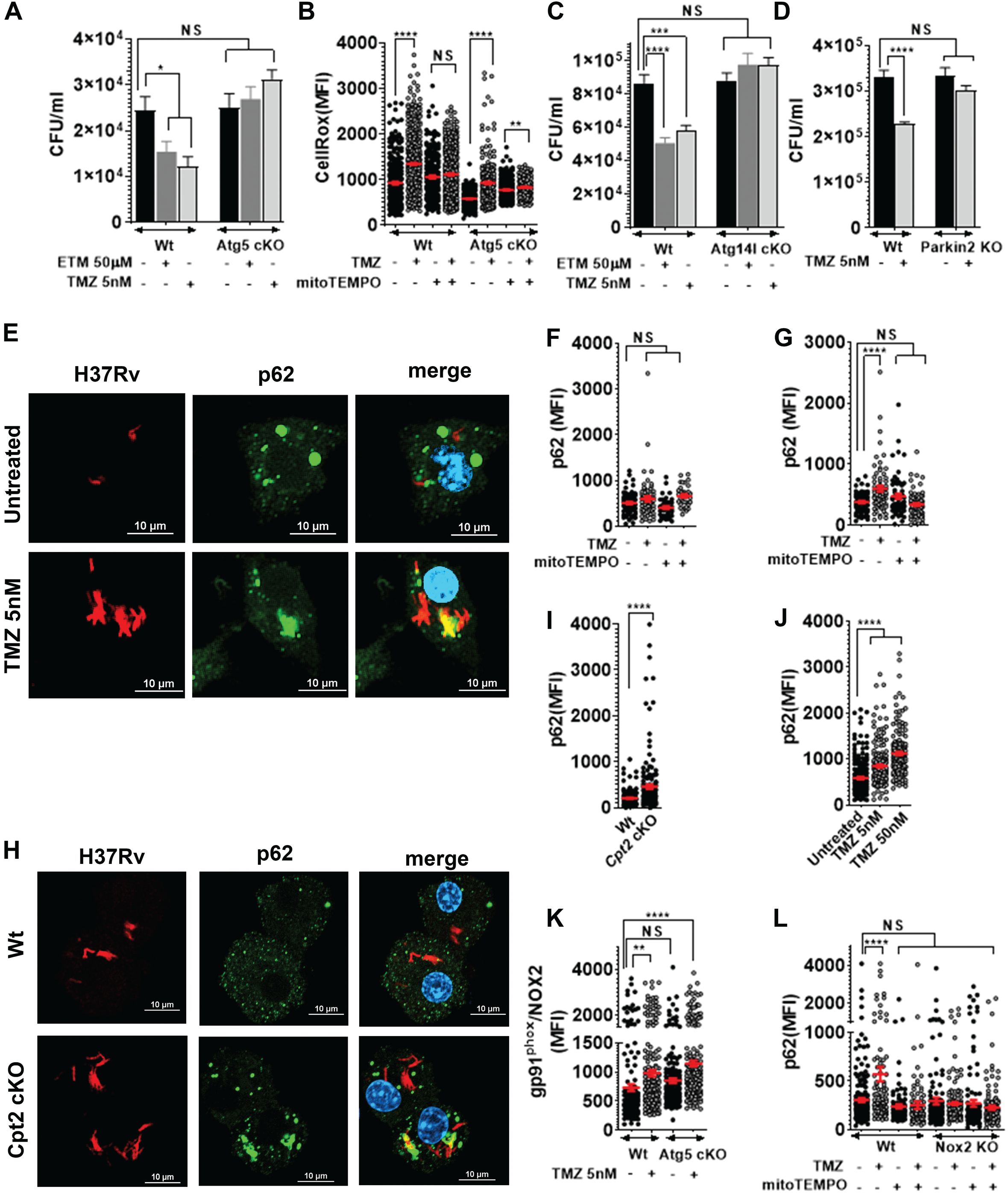
Antimicrobial activity of FAO inhibitors depends on autophagy. **(A)** Survival of Mtb was compared in FAO inhibitor treated control and *Atg5* cKO macrophages, 72hpi. **(B)** CellRox MFI in Mtb infected control and *Atg5* cKO BMDMs after TMZ treatment (5 nM, 3hr), with or without mitoTEMPO. ****P≤0.0001, unpaired Student’s t-test with Welch’s correction. Survival of Mtb in BMDMs from control and **(C)** *Atg14l* cKO cells 72 hpi, and **(D)** *Parkin2* KO cells 120 hpi. Data shows mean +/- s.e.m. *P=0.04, using unpaired t-test with Welch’s correction in (A) or ***P=0.0002, ****P≤0.0001 using ordinary one-way ANOVA in (C, D). **(E)** IF microscopy of p62 (green) and dsRed-expressing H37Rv (red) in BMDMs treated with 5nM TMZ 24hpi. The co-localized region is shown in yellow, scale bar = 10µm. MFI of p62 co-localized with Mtb was measured in BMDMs treated with 5nM TMZ alone or with mitoTEMPO for **(F)** 3 or **(G)** 24 hours. **** P≤0.0001, ordinary one-way ANOVA. **(H)** IF images and **(I)** MFI of p62 co-localized with Mtb in *Cpt2* KO and control BMDMs 24hpi. **(J)** MFI of p62 co-localized with Δ*esxA* in BMDMs treated with TMZ for 24h. **(K)** MFI of gp91^phox^/NOX2 co-localized with Mtb in control versus *Atg5* cKO BMDMs after TMZ treatment (5nM, 24hpi). **(L)** MFI of p62 co-localized with Mtb in control and Nox2KO macrophages treated with 5nM TMZ with or without mitoTEMPO, 24 hpi. Data shows mean +/- s.e.m.**P=0.001, ****P≤0.0001 calculated using unpaired Student’s t-test with Welch’s correction in (K) and ordinary one-way ANOVA for (J,L). All panels show data from one representative experiment from at least 2 independent replicates.

These studies established that FAO inhibition induced mitoROS, which promoted NADPH oxidase recruitment to the Mtb phagosome and xenophagy. Next we examined the relationship between the NADPH oxidase and xenophagy. We observed that the enhanced NADPH oxidase recruitment in response to TMZ occurred independently of autophagy, as it was also seen in *Atg5* cKO macrophages (Figure 6K). On the other hand, TMZ failed to promote xenophagy in NADPH oxidase-deficient macrophages (Figure 6L). Taken together, our data shows that mitoROS was the primary signal induced by FAO inhibition that resulted in enhanced NADPH oxidase and xenophagy, which promoted bacterial control.

## DISCUSSION

Recent work has highlighted the relationship between macrophage metabolism and inflammatory phenotype (Escoll and Buchrieser, 2019). Within the context of Mtb infection, macrophages undergo a Warburg-like shift with enhanced aerobic glycolysis (Gleeson et al., 2016; Lachmandas et al., 2016; Shi et al., 2015) and usage of TCA cycle intermediates for the generation of immuno-metabolites (Nair et al., 2018). Mtb-induced glycolysis reduces intracellular bacterial survival through enhanced IL-1*β* production, but the impact of fatty acid metabolism is not well established. Mtb also dramatically alter host lipid metabolism, promoting the formation of lipid-droplets (Cumming et al., 2018; Peyron et al., 2008a; Russell et al., 2009; Singh et al., 2012) and increasing mitochondrial dependency on exogenous fatty acids (Cumming et al., 2018). Because Mtb can utilize host fatty acids as a carbon source during infection, a prevailing idea is that host and pathogen are in competition for resources, so that reduced host consumption or enhanced production of cellular lipids would provide a nutrient source for bacterial replication (Almeida et al., 2009; Kim et al., 2017; Ouimet et al., 2016). Instead, we found that inhibition of host FAO impaired mycobacterial replication. Our initial *in vitro* results are in concordance with a recent report that showed the efficacy of FAO inhibitor ETM in Mtb-infected BMDMs (Huang et al., 2018). In that study, the antimicrobial effect was attributed to reduced IFN-β, but ETM was used at 200 μM, which has well-documented off-target effects (Divakaruni et al., 2018). We did not see effects on Type I IFN (Figure S4), and we establish the antimicrobial mechanism of FAO inhibition using specific inhibitors and genetically FAO-deficient *Cpt2* knockout macrophages. Unexpectedly, we found that blocking FAO enhanced macrophage effector functions by eliciting mitoROS, which promoted NADPH oxidase and xenophagy to restrict Mtb infection. Our results suggest mitoROS as a connecting link between macrophage fatty acid metabolism and macrophage effector functions to control Mtb infection.

The earliest cellular response that we could detect after FAO inhibition was induction of mitoROS, which occurred within three hours of treatment in both infected and uninfected macrophages. Typically, FAO products, such as NADH and FADH_2_, are oxidized by respiratory chain supercomplexes to generate the forward electron flow (FET) required for ATP production. The respiratory chain is organized into interacting supercomplexes to minimize leakage of electrons and ROS formation. Recent studies report that enzymes of FAO also physically interact with ETC components, forming an integrated, multifunctional complex (Wang et al., 2019). The FAO trifunctional protein (TFP), a tetramer that includes the target of TMZ (3-ketoacyl-CoA thiolase/HADHB), interacts with the NADH-binding domain of complex I. The presence of ETC supercomplexes dedicated to FAO may explain why FAO inhibition leads to mitoROS. In the absence of NADH shuttling from FAO to these dedicated supercomplexes, RET might occur. This fits with our finding that combined treatment with TMZ and rotenone, a complex I inhibitor, decreased mitoROS levels, consistent with RET generating the ROS at complex I. mitoROS is also generated in response to microbial products; toll-like receptor signaling alters supercomplex assembly and generates RET-ROS at complex I, both of which impact immunological signaling (Garaude et al., 2016; Langston et al., 2019; Mills et al., 2016; West et al., 2011). Thus, the mitoROS induced by TMZ may activate an antimicrobial pathway that is normally induced and protective in the context of other infection, but suboptimal in during Mtb infection. Interestingly, in distinction to our finding that FAO inhibition generated mROS, the reverse relationship was seen in *Salmonella* infected zebrafish larvae and J774.2 murine macrophage cells, where FAO enhanced *Salmonella*-induced mROS (Hall et al., 2013). The difference between this previous study and ours could be due to the experimental systems, and it may reflect a different metabolic response of macrophages to mycobacteria as compared to *Salmonella*.

The idea that FAO inhibition enhances macrophage control of Mtb by generating mitoROS is consistent with recent studies showing that mitoROS contributes to macrophage control of *Streptococcus pneumonia*, *Staphylococcus aureus*, and *Salmonella typhimurium* (Abuaita et al., 2018; Bewley et al., 2017; Hall et al., 2013; West et al., 2011). Our results suggest that mitoROS does not exert its antimicrobial function by directly killing Mtb, since mitoROS was generated in macrophages lacking *Nox2* and *Atg5*, but the antimycobacterial activity was lost. Rather, mitoROS appears to serve as a signal that enhances NADPH oxidase activity and xenophagy. A stimulatory effect of mitoROS on the NADPH oxidase has been documented most extensively in non-lymphoid cells, and recent work shows that in neutrophils mitoROS activates the NADPH oxidase and other effector functions (Nazarewicz et al., 2013; Pinegin et al., 2018; Vorobjeva et al., 2017). In addition to promoting the phagosomal recruitment of NADPH oxidase to mycobacterial phagosomes, our metabolomics data indicate that FAO inhibition may augment the PPP, which would enhance NADPH production to power the NADPH oxidase.

It will be important to understand how mitoROS promotes phagosomal assembly of the NADPH oxidase. NADPH oxidase assembly requires membrane trafficking of two integral membrane subunits (p22*^phox^* and gp91*^phox^*) followed by recruitment of the cytosolic subunits (p40*^phox^*, p47*^phox^*, p67*^phox^*). Normally this occurs shortly after uptake of phagocytic cargo that activate pathogen recognition receptors. In our experimental system, the kinetics are very different than what is typically described. We added TMZ 4h after Mtb had been added to macrophages and when extracellular bacilli had been washed away. We found that TMZ enhanced localization of both the gp91*^phox^* and p40 *^phox^* subunits with mycobacterial phagosomes 24 hours later. This suggests that signals generated by mitoROS either overcome a Mtb-induced blockade to NADPH oxidase recruitment or promote recruitment through a previously unappreciated pathway. How mitoROS regulates phagosomal NADPH oxidase recruitment and assembly will be important to establish in future studies, as our work indicates that it can be augmented, even after bacterial uptake, to promote microbial clearance of intracellular bacilli. This might be a way to clear foci of persistent bacilli.

In addition to NADPH oxidase recruitment, mitoROS also promoted xenophagy. Xenophagy and LAP are related LC3-trafficking pathways that can promote microbial clearance (Upadhyay and Philips, 2019). They also differ, as LAP occurs immediately after phagocytosis and involves LC3 recruitment to a single phagosomal membrane, whereas xenophagy depends upon formation of a double membrane compartment. We found that the antimicrobial activity of TMZ depended on ATG14L and PARKIN, which are required for canonical autophagy and Mtb xenophagy (Manzanillo et al., 2013). In addition, TMZ enhanced the association of phagosomes with p62, a selective autophagy adaptor, and the response occurred well after bacterial uptake, all of which suggest that the mechanism of clearance is xenophagy and not LAP, despite the involvement of NADPH oxidase which has been linked to LAP. We also cannot rule out that both LAP and xenophagy are enhanced by FAO inhibition. In terms of how the NADPH oxidase might activate xenophagy, one possibility is that the NADPH oxidase generates phagosomal membrane damage, which is known to promote pathogen ubiquitination, adaptor recruitment, and xenophagy (Manzanillo et al., 2012; Wong and Jacobs, 2011). We observed NADPH oxidase recruitment in TMZ-treated *Atg5* cKO macrophages, yet, TMZ failed to kill Mtb in the absence of xenophagy, arguing against direct antimycobacterial activity of the NADPH oxidase. The lack of direct activity is consistent with other recent studies (Köster et al., 2017; Olive et al., 2018), and likely reflects that Mtb has ROS-detoxifying activities such as KatG, a catalase/peroxidase. We conclude that the antimycobacterial activity of ROS is primarily based upon its ability to activate lysosomal trafficking pathways, perhaps due to damage to the phagosomal membrane resulting in xenophagy activation.

While, the current view is that ROS generation contributes to inflammation and cellular damage, a beneficial role of RET-ROS was appreciated recently for hypoxia sensing in the carotid body (Fernández-Agüera et al., 2015), ischemia-reperfusion injury (Chouchani et al., 2014), and extending fly lifespan (Scialò et al., 2016). A number of studies propose the use of FAO inhibitors as metabolic therapy in cancer (Duman et al., 2019; Wang et al., 2018) and heart failure (Lionetti et al., 2011). Given the link that we have shown between FAO inhibition and mitoROS, it will be important to establish whether that contributes to their therapeutic effects. TMZ, which has been used to treat angina for 35 years, is orally bioavailable, lacks substantial drug-drug interactions, and has a favorable safety profile (Dézsi, 2016). Importantly, TMZ showed antimycobacterial activity in mice at drug concentrations that are used clinically. Therefore, TMZ could be rapidly translated into an HDT to be used as an adjunct to conventional antibiotics. The magnitude of Mtb control was modest, but similar to other HDTs that have been tested in mouse models of Mtb (Napier et al., 2011; Parihar et al., 2014; Singhal et al., 2014). In the case of TMZ, the limited activity is probably not due to inadequate drug exposure, but may reflect that TMZ works in only certain myeloid cells *in vivo*, perhaps based upon their metabolic phenotype, or that there is limited capacity of mitoROS or autophagy to activate robust clearance. Distinguishing these possibilities will be important in terms of designing effective HDTs for clinical use. Evaluating the effect of TMZ on other cell types critical for Mtb infection, including neutrophils, dendritic cells, and T cells will be important. Many of the leading candidates for HDT, such as metformin and sirtuin-1, also promote autophagy (Cheng et al., 2017; Singhal et al., 2014), and whether enhanced antimicrobial control could be established by a combination of HDTs that target distinct cellular reservoirs or work synergistically to activate antimicrobial pathways will be critical to investigate.

## ACKNOWLEDGEMENTS

We thank members of the Philips laboratory and Robert Mahon (NIH/NIAID) for critical reading of the manuscript. We thank Ronald E. Dolle (Washington University in St. Louis (WUSTL)) for help designing PK studies, Michael J. Wolfgang (Johns Hopkins University School of Medicine) for providing Cpt2*^fl/fl^* mice, and Jeff Cox (University of California, Berkeley) for providing Rv-lux strain. We thank the Alvin J. Siteman Cancer Center (WUSTL), Barnes-Jewish Hospital in St Louis, MO, the Bursky Center for Human Immunology and Immunotherapy Programs Immunomonitoring Laboratory, and Diane Bender for the multiplexing immunoassay service. The Siteman Cancer Center is supported in part by an NCI Cancer Center Support Grant #P30 CA091842. This work was supported by grants from the NIH (R21 AI128427), the Center for Drug Discovery (WUSTL), and LEAP Inventor Challenge Award (WUSTL) to J.A.P, NIH (R35 HL135799) to K.J.M., NIH (R01 DK11003404) to J.D.S., and the Pott’s Memorial Foundation to P.C.

## AUTHOR CONTRIBUTIONS

M.O. and S.K. did experiments demonstrating activity of etomoxir against intracellular Mtb and P.C. performed all subsequent experiments. She had help with mice and MIC measurements from G. Y. and SeaHorse experiments by L.H. M.Z., H.W., and V.D. measured TMZ concentrations from serum of infected mice. J.A.P. conceived the project and supervised the experiments with input from J.D.S. P.C. and J.A.P. wrote the manuscript; P.C., S.K., M.O., K.J.M., V.D., J.D.S, and J.A.P. edited the manuscript.

## STAR Methods

### KEY RESOURCES TABLE

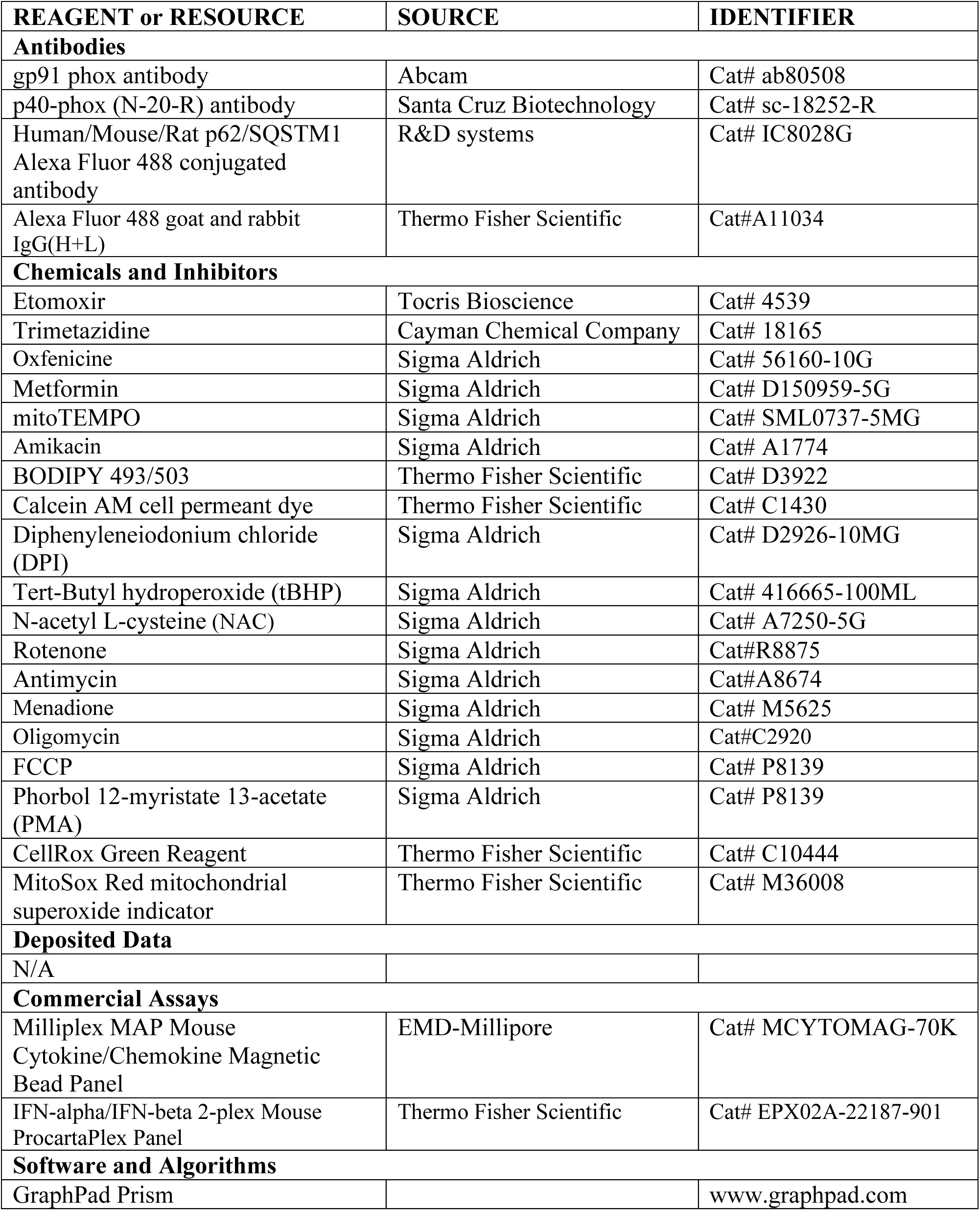
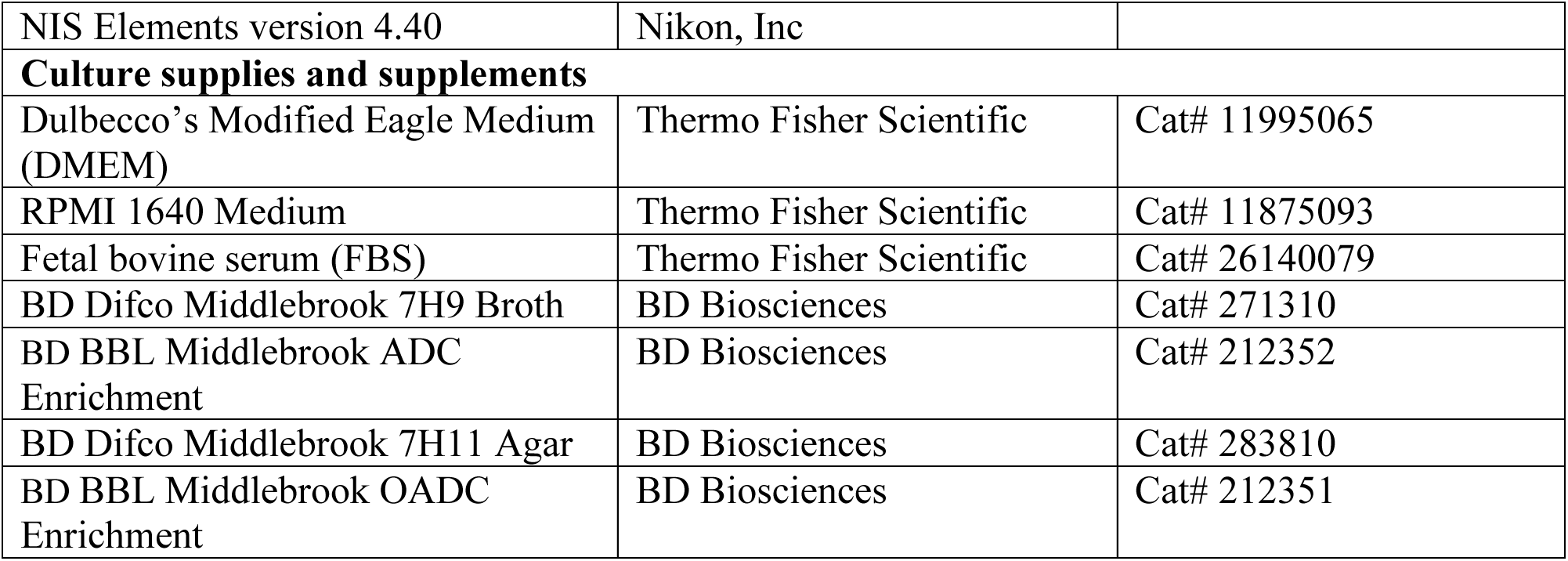

## LEAD CONTACT AND MATERIALS AVAILABILITY

Further information and requests for reagents may be directed to and will be fulfilled by the Lead Contact, J. A. Philips (philips.j.a<u>@wustl.edu)</u>

## EXPERIMENTAL MODEL DETAILS

### Bacterial Strains

*M. tuberculosis* and *M. abscessus* were grown at 37°C to log phase in Middlebrook 7H9 broth (BD Biosciences) supplemented with 0.05% Tween 80, BD BBL Middlebrook ADC Enrichment (BD Biosciences), and 0.2% (vol/vol) glycerol. Plasmids were selected with 25 µg/ml kanamycin or 50 µg/ml hygromycin depending on the resistance marker. H37Rv, the wild type Mtb strain, and Δ*esxA* were provided by William Jacobs Jr. (Albert Einstein College of Medicine)(Wong and Jacobs, 2011). DsRed expressing H37Rv was a gift from J. Ernst (New York University). H37Rv expressing the *Vibrio harvei* luciferase (Rv-lux) was gift from Jeff Cox (University of California, Berkeley). The “live-dead” reporter strain constitutively expresses mCherry and inducibly express GFP under control of a tetracycline-inducible promoter (Martin et al., 2012).

### Mice

We used 8-12 week old C57BL/6 mice. *Cpt2^fl/fl^* LysM-Cre^+^ mice were generated by crossing *Cpt2^fl/fl^* and LysM-Cre^+^ mice. The generation of *Cpt2^fl/fl^* and LysM-Cre^+^ mice have been described previously (Clausen et al., 1999; Lee et al., 2015). The Washington University School of Medicine Institutional Animal Care and Use Committee approved all the work with mice. Euthanasia was performed prior to bone marrow harvest in accordance with the 2013 *AVMA Guidelines for the Euthansia of Animals* (https://www.avma.org/KB/Policies/Documents/euthanasia.pdf).

## METHOD DETAILS

### Cell culture

To obtain murine bone marrow-derived macrophages (BMDMs), marrow was flushed from the femurs and tibia of mice, and the hematopoetic stem cells were allowed to differentiate for 7 days in Dulbecco’s Modified Eagle Medium (DMEM, Gibco) supplemented with 10% fetal bovine serum (FBS, Gibco), 1% Pen-Strep solution (Gibco), and 20% L929-conditioned medium. After 7 days, the BMDMs were harvested using Ca^2+^/Mg^2+^-free PBS (Gibco) containing 5mM EDTA (Invitrogen, Life Technologies), and maintained in DMEM containing 10% FBS and 10% L929-conditioned medium after infection. Immortalized BMDMs (iBMDMs) were immortalized by infection with the J2 retrovirus (BEI Resources). RAW 264.7 and THP-1 cells were obtained from American Type Tissue Collection (ATCC) and were maintained in DMEM and RPMI-1640, respectively, with 10%FBS. THP-1 differentiation was induced using 20 ng/ml phorbol-12-myristate acetate (PMA, Sigma) for 18-20 hours.

### Bacterial infections

For *in vitro* macrophage assays, a log phase culture of Mtb H37Rv was pelleted and resuspended in macrophage culture medium. Bacterial single-cell suspensions were prepared by filtering through 5-micron filters (PALL Life Sciences, catalog no. 4650). The number of Mtb in the resultant filtrate was estimated by measuring absorbance at 600 nm, followed by infection of macrophages at a multiplicity of infection (MOI) of 5. After 4 hours, macrophages were washed three times with warm DMEM to remove extracellular bacteria, and then resuspended in medium containing FAO inhibitors or solvent control. To estimate intracellular Mtb growth, infected macrophages were lysed in 0.06% SDS solution at the indicated time points, and serial dilutions of the lysates were plated on 7H11 agar plates (BD Biosciences, catalog no. 283810) containing BD BBL Middlebrook OADC Enrichment (BD Biosciences, catalog no. 212351) and glycerol. Colony forming units (CFU) were calculated 14-21 days later. For *in vivo* infections, log phase H37Rv culture was pelleted and resuspended in sterile 0.5% Tween 80 solution. After a centrifugation step, the supernatant was used for aerosol infection using a Glas-Col inhalation exposure system. The infectious dose administered was calculated by plating CFU from an aliquot of the bacterial suspension.

### Mice surgeries and aerosol infections

Alzet mini-osmotic pumps (Model 2002, Durect Corporation, CA) were loaded with saline or TMZ solution as per the manufacturer’s protocol. The osmotic pumps were surgically implanted in anesthetized mice under aseptic conditions. The mice were administered analgesic to minimize pain and were monitored regularly for signs of pain and other postoperative complications. Two days later, the mice were infected with H37Rv via aerosol route using an inhalation exposure system from Glas-Col. The dose of infection was confirmed one day post-infection by plating whole lung homogenates from 2 mice on Middlebrook 7H11 agar. 2 weeks post infection, the mice were euthanized and the lungs were harvested, homogenized, and plated for CFUs.

### Serum microsampling of TMZ

Serum concentrations of TMZ in mice were determined at 5 and 14 days after initiating treatment. Lidocaine was applied to mice tails to minimize pain, and the end of the tail was wiped with alcohol and a small incision was made. 100 µl blood was collected in K_2_-EDTA microvette tubes (Braintree Scientific, Inc) and was centrifuged at 5,000 rpm for 5 minutes to recover plasma. Samples were stored at – 80°C until analyzed for TMZ content.

### High pressure liquid chromatography coupled to tandem mass spectrometry (LC/MS-MS) analytical method

1 mg/mL DMSO stocks of Trimetazidine (Sigma-Aldrich) were serial diluted in 50/50 Acetonitrile water and subsequently serial diluted in drug free CD1 mouse plasma (K_2_EDTA, Bioreclamation IVT, NY) to create standard curves and quality control (QC) spiking solutions. Standards, QC, controls, and study samples were extracted by combining 10 µL plasma with 100 µL acetonitrile/methanol 50/50 containing the internal standard verapamil. Extracts were vortexed and centrifuged, and supernatant was transferred for LC-MS/MS analysis. LC/MS-MS was performed on a Sciex Applied Biosystems Qtrap 6500+ triple-quadrupole mass spectrometer coupled to a Shimadzu 30ACMP HPLC system, and chromatography was performed on an Agilent Zorbax XDB-C_18_ column (3×75 mm; particle size, 3.5 µm). Milli-Q deionized water with 0.1% formic acid was used for the aqueous mobile phase and 0.1% formic acid in acetonitrile for the organic mobile phase. Multiple-reaction monitoring (MRM) of parent/daughter transitions in electrospray positive-ionization mode was used to quantify all molecules. MRM transitions of 267.08/181.10 and 455.40/165.20 were used for TMZ and Verapamil respectively.

### Fluorescence microscopy and image analyses

BMDMs were seeded in 8-well chamber slides (Falcon culture slide 8-well, catalog no. 08-774-26), and infected with DsRed-expressing H37Rv at MOI 5. At the indicated time points, samples were fixed overnight with 1% paraformaldehyde (PFA). For immunofluorescence (IF), samples were permeabilized and blocked in PBS with 0.05% saponin and 3% BSA, and stained with the indicated primary antibodies for 2 h at room temperature or overnight at 4°C. Primary antibodies used were p40^phox^ (Santa Cruz Biotechnologies), gp91^phox^/NOX2 (abcam), and p62 (R&D systems). Staining with alexa fluorophore-conjugated secondary antibody (Molecular Probes) was done for 2 h at room temperature. Following this, the samples were washed with 0.1% Tween 20/PBS and mounted using Prolong Gold antifade (Thermo Fisher Scientific, catalog no. P36930). Images were captured using a Nikon Eclipse Ti confocal microscope (Nikon Instruments Inc.) equipped with a 60X apochromat oil-objective lens, and analyzed using NIS-Elements version 4.40 (Nikon). Briefly, a region of interest (ROI) was drawn around each bacterium and the mean fluorescence intensity (MFI) was measured using the ROI statistics tool.

### ROS measurement assays

For ROS measurement assays, the macrophages were seeded in 96 well plates (µ-Plate 96 well, IBIDI catalog no. 89626). To estimate total cell ROS, the samples were treated with Cell Rox green fluorescent dye (Thermo Fisher Scientific) at 5 µM for 30 minutes. The samples were washed three times with PBS, and fixed overnight with 1% PFA. Mitochondrial ROS was measured in live, uninfected macrophages using MitoSox fluorescent dye (Thermo Fisher Scientific), at 5µM for 30 minutes. The samples were imaged using confocal microscope, and the ROS signal of each cell was quantified using NIS-Elements software. Briefly, each cell was converted to a ROI and the MFI was measured using the ROI statistics tool.

### Oxygen consumption rate and extracellular acidification rate

BMDMs from *Cpt2* cKO mice and control littermates were plated in 96 well Seahorse plates at a density of 75,000 cells per well. The cells were treated with 5nM TMZ or solvent control for 3 h. After treatment, the cells were washed and placed in XF media (non-buffered RPMI 1640 containing 25mM glucose, 2mM L-glutamine and 1mM sodium pyruvate) with 10 % FCS. Oxygen consumption rate (OCR) and extracellular acidification rates (ECAR) were measured under basal conditions and following inhibitors were added: 1µM oligomycin (Sigma), 1.5µM fluorocarbonyl cyanide phenylhydrazone (FCCP, Sigma), and 100 nM rotenone (Sigma) + 1µM antimycin A (Sigma). Measurements were taken using a 96 well Extracellular Flux Analyzer (Seahorse Bioscience, North Bellerica, MA, USA).

### Metabolomics

For metabolic profiling of murine BMDMs, 15 million macrophages were seeded in 15 cm petri dishes (Genesee Scientific) with 5 replicates for each condition. The next day, macrophages were infected with H37Rv at an MOI 5 for 4 h as described above. The samples were washed and maintained in culture medium with or without with 50 nM TMZ for 3 or 24 h. At the respective time points, the samples were washed twice with sterile Hank’s Balanced Salt Solution (HBSS, Gibco), and the metabolites were extracted in 80% methanol (Sigma) in water (Corning). The samples were stored at −80° C and shipped to Metabolon Inc., NC (www.metabolon.com) for further processing and analyses.

At Metabolon, the samples were prepared using the automated MicroLab STAR^®^ system from Hamilton Company. After addition of recovery standards and protein removal, the extracts were divided into fractions for analysis by: two separate reverse phase (RP)/ UPLC-MS/MS with positive ion mode electrospray ionization (ESI), RP/UPLC-MS/MS with negative ion mode (ESI), and HILIC/UPLC-MS/MS with negative ion mode ESI. All methods utilized a Waters ACQUITY ultra-performance liquid chromatography (UPLC) and a Thermo Scientific Q-Exactive high resolution/accurate mass spectrometer interfaced with a heated electrospray ionization (HESI-II) source and Orbitrap mass analyzer operated at 35,000 mass resolution.

### Cytokine measurements

Culture supernatants were harvested from uninfected or Mtb-infected BMDMs that were untreated or treated with 5 or 50 nM TMZ for 24 and 72 h. The conditioned media was filter-sterilized and cytokines and chemokines were measured using Milliplex MAP Mouse Cytokine/Chemokine Magnetic Bead Panel (MCYTOMAG-70K, Millipore Sigma) and Procartaplex Mouse IFNα/IFNβ Panel (2plex) (ThermoFisher Scientific).

### Liquid MIC determinations

In 96 well plates, Mtb cultures with starting OD_600_ of 0.01, 0.05, and 0.10 were incubated with increasing concentrations of drugs in triplicate. The growth of Mtb was measured at day 0, 1, 2, 3 and 4. The MIC was considered the minimal concentration tested that inhibited Mtb growth at 4 days. To assess the direct toxicity of TMZ on Mtb, H37Rv expressing the *Vibrio harvei* luciferase (Rv-lux) was cultured in 7H9 broth with or without 1 mM TMZ for 7 days. Relative luminescence units (RLU) were measured every 2 days at 490 nm.

### PK studies

PK and plasma binding studies were performed by Alliance Pharma (PA, USA) following IV and oral administration of 3 and 15 mg/kg TMZ, respectively. PK study of TMZ using Alzet osmotic pumps was conducted by Paraza Pharma Inc, Canada. Briefly, Alzet osmotic pumps (model 2002, Durect Corporation) were surgically implanted in C57BL/6 mice (n=5) for a subcutaneous infusion at 10.66 m/k/day. Plasma TMZ concentrations were determined over the course of 48h.

### Statistics

GraphPad Prism software was used to prepare plots and assess statistical significance of results using unpaired Student’s t-test with Welch’s correction, Ordinary one-way ANOVA and Mann-Whitney test.

## SUPPLEMENTAL INFORMATION

**Figure S1.**
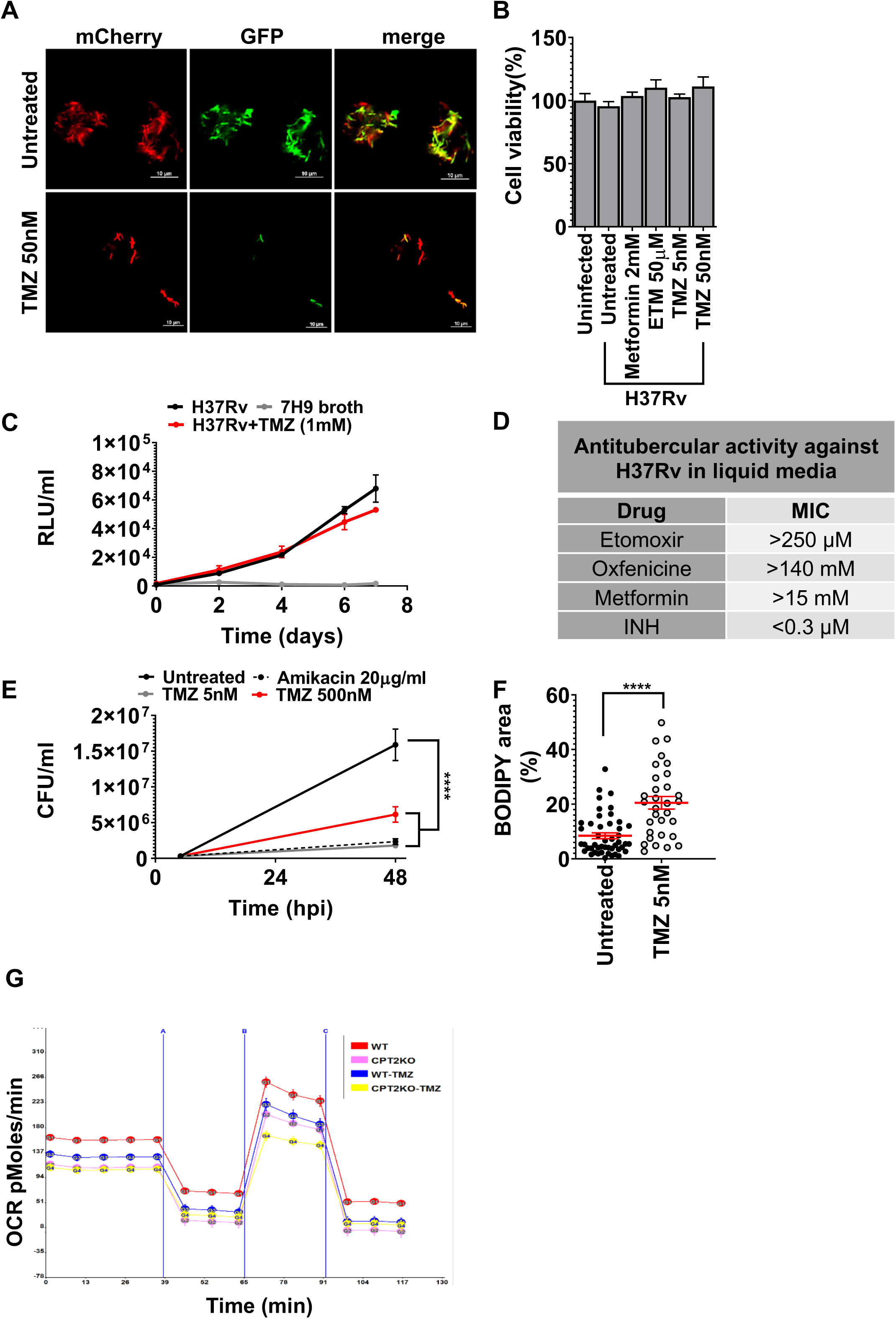
Inhibition of macrophage FAO restricts intracellular *M. tuberculosis* growth. **(A)** Murine BMDMs were infected with live-dead reporter expressing H37Rv and were either untreated or treated with 50 nM TMZ for 6 days. The reporter Mtb strain constitutively expresses mCherry and has GFP controlled by a tetracycline-inducible promoter. To induce GFP expression, 200 nM anhydrotetracycline was added 2 days after infection. Metabolically active bacteria express both GFP and mCherry, whereas dead bacteria are only mCherry-positive. Representative images show more dead Mtb upon TMZ treatment as compared to control. Scale bar= 10 µm. **(B)** Macrophage were uninfected or infected with Mtb for 72h, and cell viability was assessed using calcein dye mean fluorescence intensity (MFI). Plot shows cell viability expressed as % of uninfected. Data shows mean +/- s.e.m., *P=0.03, **P≤0.0005 ordinary one-way ANOVA. (**C)** H37Rv expressing the *Vibrio harvei* luciferase (Rv-lux) was cultured in 7H9 broth with or without 1 mM TMZ for 7 days. Plot shows bacterial growth as assessed by relative luminescence units (RLU). (**D)** Minimal inhibitory concentration (MIC) of etomoxir, oxfenicine, metformin, and isoniazid (INH, positive control) was determined by culturing H37Rv for 4 days in 7H9 broth supplemented with various doses of inhibitors and measuring absorbance at 600 nm. (**E)** Intracellular growth of *M. abscessus* in BMDMs that were untreated or treated with indicated concentrations of amikacin or TMZ for 48 hours. Data shows mean +/- s.e.m., ****P≤0.0001 ordinary one-way ANOVA. (**F)** BODIPY fluorescent dye was used to stain lipid bodies in BMDMs that were infected for 24 hours with Mtb and either untreated or treated with TMZ. Shown here is mean +/- s.e.m of the % area in a cell occupied by BODIPY stain. ****P≤0.0001, unpaired Student’s t-test with Welch’s correction. Using extracellular flux analysis, we quantified the (**G)** oxygen consumption rate (OCR) in uninfected control and *Cpt2* cKO BMDMs treated with 5nM TMZ or solvent control for 3h, followed by sequential addition of oligomycin (A), FCCP (B), and rotenone plus antimycin (C). Data shows average +/- s.d. of 16 replicates.

**Figure S2.**
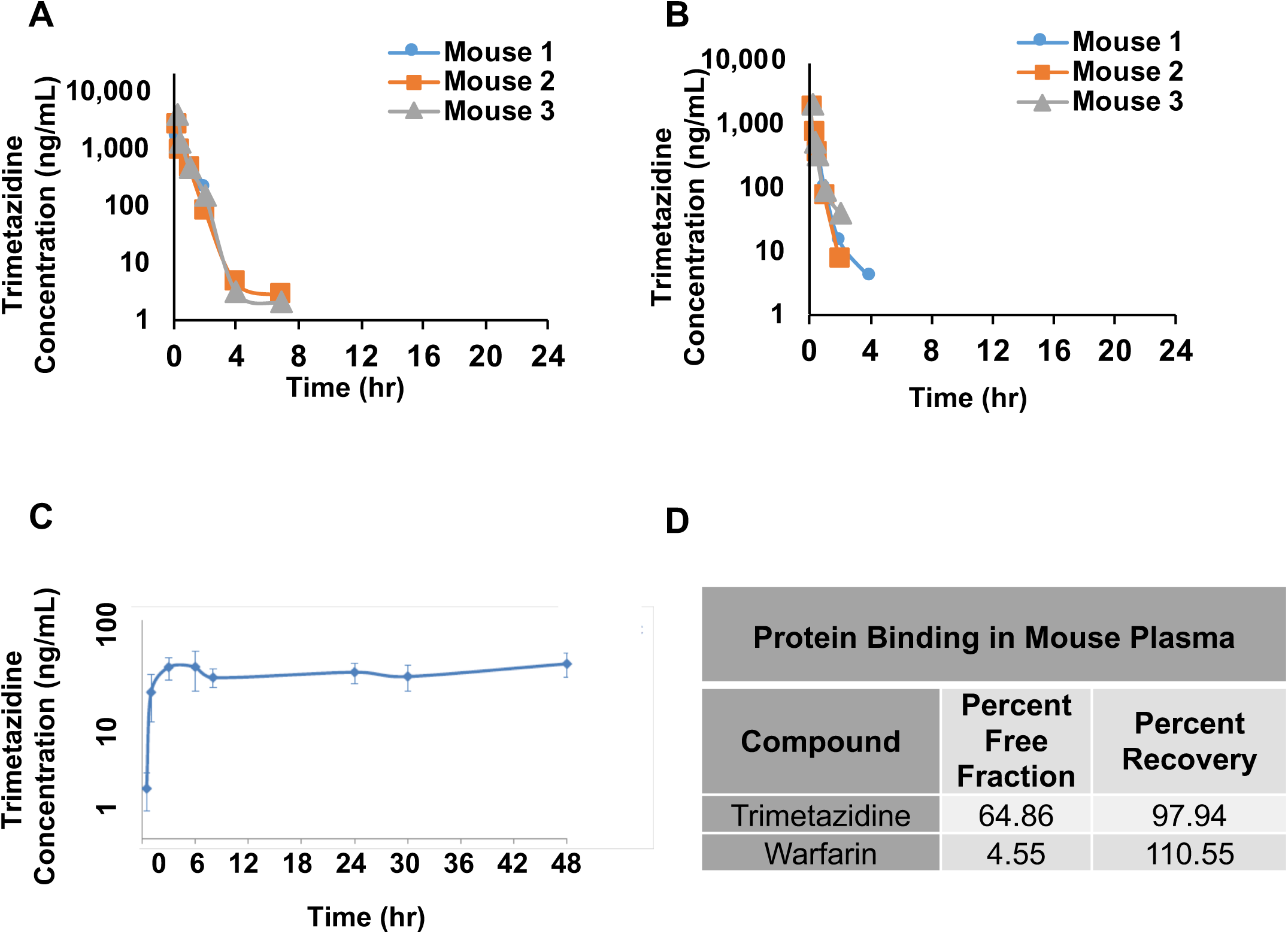
Pharmacokinetics study of TMZ in mice. The half-life of TMZ was determined in C57BL/6 mice that were administered the inhibitor **(A)** orally at 15 mg/kg or **(B)** intravenously at 3 mg/kg. **(C)** TMZ (10.66 mg/kg/day) was administered to female C57Bl/6 mice (n=5) for 48 hours using Alzet osmotic pumps and achieved C_ave_-34.5 ng/ml. **(D)** Protein binding of TMZ in mouse plasma (C57BL/6) was measured using rapid equilibrium dialysis (n=3). Warfarin was used as a control. In humans TMZ is reportedly weakly protein bound (∼16%)(Harpey et al., 1988).

**Figure S3.**
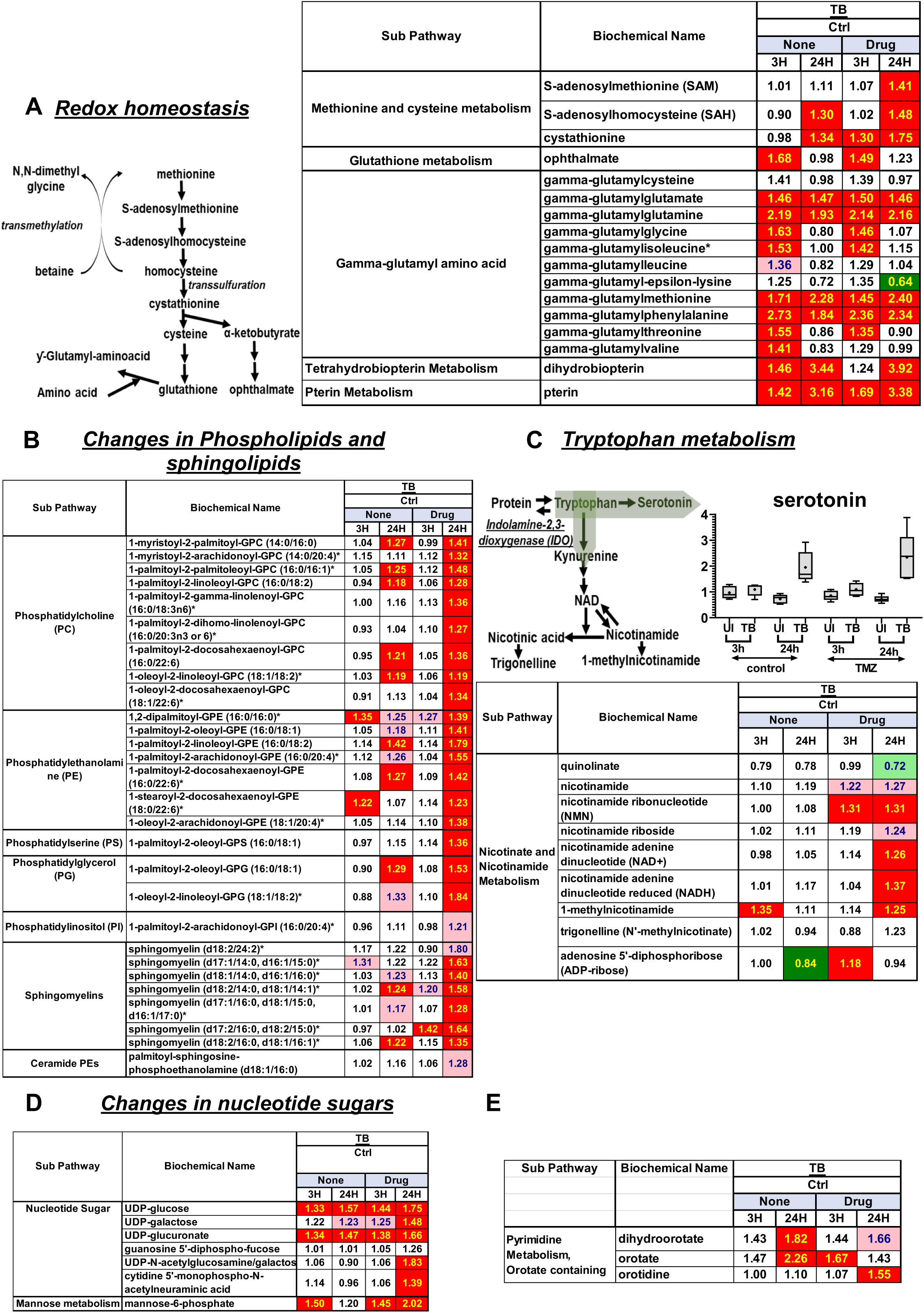
FAO inhibition enhances metabolic changes in response to Mtb infection. Unbiased metabolite profiling was performed in BMDMs to characterize the effect of Mtb infection and TMZ treatment. **(A)** Macrophage response to infection involved perturbations in redox homeostasis. Increased glutathione synthesis was suggested by increased methionine to cysteine conversion pathway metabolites SAM, SAH and cystathione. Ophthalmate, a compositional derivative of glutathione, can reflect enhanced activity of glutathione synthetase. Marked elevations were observed in gamma-glutamyl amino acids, which are formed when gamma-glutamyl transpeptidase transfers the gamma-glutamyl moiety of glutathione to acceptor amino acids. Infection significantly increased dihydrobiopterin and biopterin. These metabolites are oxidized forms of tetrahydrobiopterin, which is a cofactor for all NOS isoforms and its depletion is associated with enzyme uncoupling and ROS generation. **(B)** Increases in membrane phospholipids and sphingomyelin were detected 24 hpi, which were enhanced in presence of TMZ. **(C)** Tryptophan metabolism was increased with infection 24 hpi. Elevated levels of serotonin were observed, with more pronounced accumulation upon TMZ treatment. Graph shows ScaledImpData of serotonin in uninfected (UI) and infected (TB) samples at the indicated time points. The line and dot show median and mean, respectively. Tryptophan can be converted by indoleamine 2,3-dioxygenase (IDO) to kynurenine, which is further utilized for nicotinamide metabolism. TMZ treatment in infected cells increased levels of metabolites in the nicotinamide pathway, suggesting it might improve infection control in combination with SIRT 1 activators, as sirtuins are NAD+-dependent class III histone deacetylases and activating sirtuin 1 (SIRT 1) is protective in Mtb infection (Cheng et al., 2017). Other metabolic changes during infection included (**D**) increase in nucleotide sugars, with a marked increase in UDP-N-acetylglucosamine/galactosamine (UDP-GlcNAc) when FAO was inhibited. These metabolites are utilized in protein and lipid glycosylation reactions, and UDP-GlcNAc biosynthesis was shown to be important for M2 macrophage polarization (Jha et al., 2015). (**E**) TMZ-induced accumulation in uracil-containing pyrimidines in Mtb infected macrophages. This may indicate an increase in dihydroorotate dehydrogenase (DHODH) activity, which catalyzes a rate-limiting step in the pyrimidine pathway. Since DHODH can supply electrons downstream of complex I of the mitochondrial ETC, it can potentially contribute to RET-ROS production. (A-F) Heatmaps show metabolite ratios in Mtb infected and uninfected (TB/Ctrl) macrophages, that were untreated (None) or TMZ treated (Drug) for 3 or 24 h. Here, significant difference (p≤0.05, three-way ANOVA) between groups are colored in green for metabolite ratio of <1 and red for ratio of >1. Light green and red colors show groups that missed the statistical cutoff for significance 0.05<p<0.10.)

**Figure S4.**
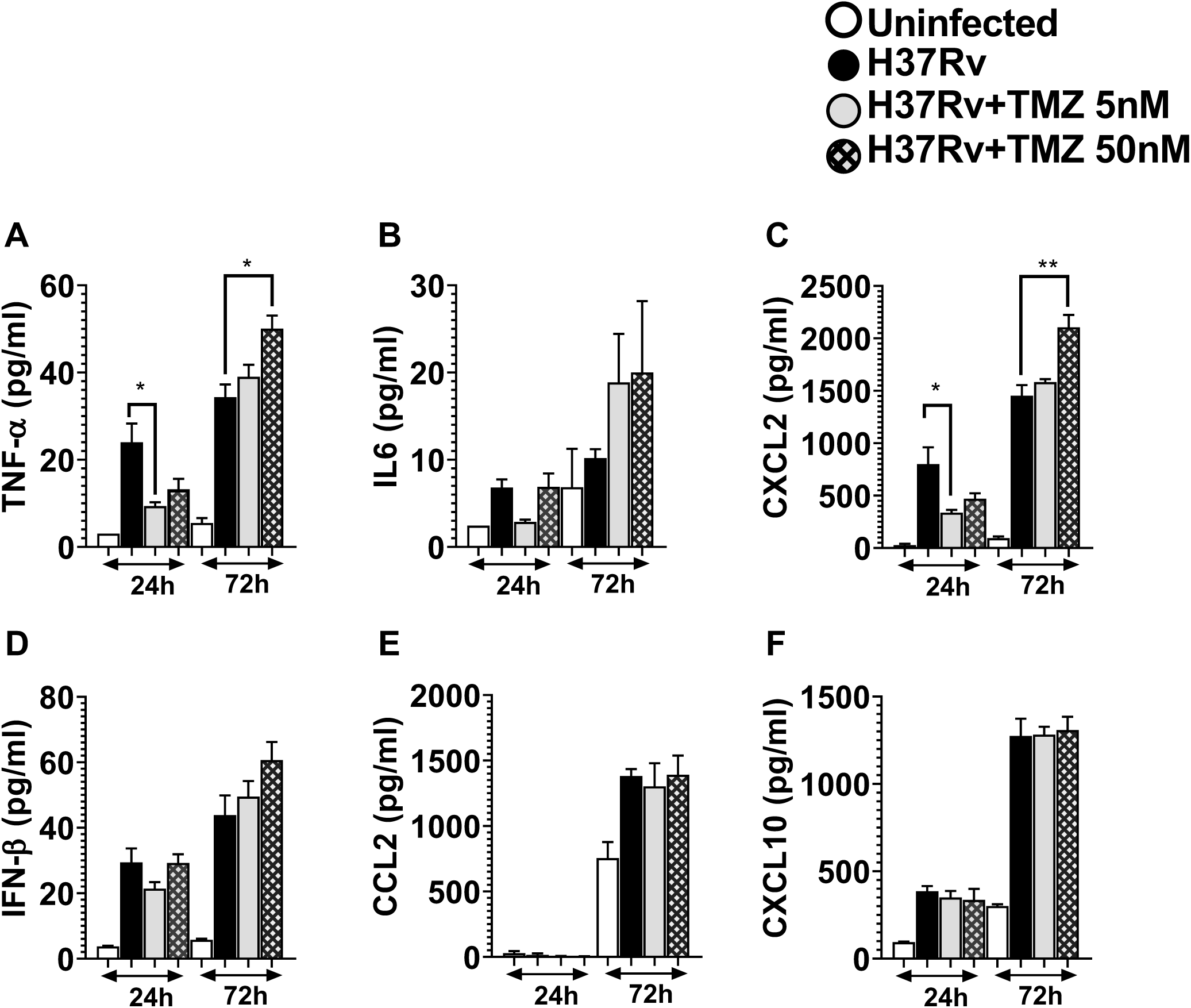
Pro-inflammatory cytokine secretion in macrophages treated with TMZ. Supernatants from uninfected or Mtb-infected BMDMs that were untreated or treated with indicated concentrations of TMZ were harvested 24 and 72 hpi. The levels of pro-inflammatory cytokines and chemokines **(A)** TNF-α **(B)** IL-6, **(C)** CXCL2, **(D)** IFN-β, **(E)** CCL2, and **(F)** CXCL10 were measured using a Milliplex MAP Mouse Cytokine/Chemokine Magnetic Bead Panel. *P≤0.02, **P=0.004 calculated using ordinary one-way ANOVA.

**Table S1. Metabolite Differences Associated with Mtb infection and TMZ treatment**

